# Independent Basal Ganglia Neural Populations Encode Speech Production and Ambient-Noise Levels

**DOI:** 10.1101/2025.11.18.689157

**Authors:** Latané Bullock, Matteo Vissani, Alan Bush, Jackie S. Kim, Lori L. Holt, Julie Fiez, Robert S. Turner, Todd M. Herrington, Jeffrey Schweitzer, Frank H. Guenther, R. Mark Richardson

## Abstract

The function of the basal ganglia (BG), a set of nuclei deep in the brain, is still under debate despite their clinical relevance. Lines of evidence from non-human primate and human experiments suggest that one key causal contribution of the adult primate BG is invigoration of movement according to environmental and energetic constraints. Previous studies have come to this conclusion using primarily limb motor control tasks. We hypothesized BG are also important for invigorating the respiratory, vocal, and articulatory motor systems during speech production. To test this, we used a rare clinical opportunity to record local field potentials-broadband gamma activity (LFP-BGA) and single unit (SU) activity from human basal ganglia during deep brain stimulation implant surgeries. Participants repeated sentences aloud in the operating room. We modulated environmental and energetic constraints of speech production by inducing the Lombard Effect, wherein people reliably increase the intensity of their voice in the presence of background noise. Participants increased voice intensity, pitch, vowel space size, and respiratory depth in the presence of ambient noise. ∼40% of LFP and SU sites were modulated by speech production. Comparing globus pallidus (GP) LFP-BGA, herein reported for the first time during speech, to subthalamic nucleus (STN) LFP-BGA revealed GP activated after STN. STN was more coincident with the speech preparatory period and GP more coincident with speech onset, suggesting differential roles in speech production. We found only weak evidence that STN or GP activity encodes the level of produced speech intensity. Instead, we found that LFP-BGA and SU-FR robustly track the ambient background noise levels. Linear decoding from SU-firing rate revealed we could infer the background noise levels with >75% accuracy in any given time point in the trial, including baseline. Importantly, the ambient-noise-modulated sites were an independent population from speech-modulated sites. We conclude that human BG encode the energetic constraints of communication. We theorize that the code is leveraged in downstream BG-cortical loop nodes to invigorate speech oro-motor gestures and co-speech manual limb gestures, both of which respond to ambient noise levels to maintain high-fidelity communication.

## Introduction

Humans seamlessly change voice intensity according to communicative and energetic constraints; we speak quietly in quiet spaces, and loudly in the presence of background noise to ensure our interlocutors receive our intended signals with high fidelity (Brumm & Zollinger, 2011; Lombard, 1911; Luo et al., 2017). This phenomenon was named the ‘Lombard effect’ after French otolaryngologist Étienne Lombard, who noted that patients in hospital environments increased their voice intensity in response to ambient noise (Lombard, 1911; Luo et al., 2018). The Lombard effect has since been described in many vocal vertebrates, including birds, whales, cats, bats, and non-human primates (Brumm & Zollinger, 2013; Hotchkin & Parks, 2013; Pedersen et al., 2024). The effect across species is best predicted by the frequency-specific signal-to-noise ratio (SNR) of the vocalization relative to ambient noise, rather than any background noise per se. Researchers have noted that, in humans, the Lombard effect is not simply indexed by an increase in vocal intensity but several other speech properties, including changes in fundamental frequency, vowel duration, and the size of a speaker’s vowel space (Stowe & Golob, 2013). This constellation of speech-in-noise changes is sometimes called ‘Lombard speech’.

Despite the scientific and clinical relevance of the Lombard effect (Adams & Lang, 1992; Stathopoulos et al., 2014), its underlying neural substrates remain unclear. The cross-species Lombard effect is likely mediated by subcortical circuits (Luo et al., 2018; Nonaka et al., 1997), but Lombard speech in humans is modulated by social context and complex cognitive factors that may engage the whole speech production system (Alghamdi et al., 2018; Stowe & Golob, 2013; Trujillo et al., 2021). Clinical observations and studies using animal models of non-speech motor control suggest that proxies of motor ‘vigor’—like movement velocity, amplitude, or frequency—are encoded in the basal ganglia (BG), a set of functionally connected nuclei deep in the brain. This idea is supported by studies of non-invasive human neuroimaging (Pope et al., 2005; Spraker et al., 2007), clinical symptomology in diseases of the basal ganglia (Ramig et al., 2008; Sapir, 2014; Viviani et al., 2009), non-human primate (NHP) globus pallidus internus (GPi) lesions (Desmurget & Turner, 2008, 2010; Horak & Anderson, 1984), NHP single neuron recordings from BG (Mink & Thach, 1991a, 1991b; Thura & Cisek, 2017; Turner & Anderson, 1997), and human intraoperative limb movement recordings (Brücke et al., 2008, 2012; Fischer et al., 2017; Jenkinson et al., 2013; Joundi et al., 2012; Nonaka et al., 1997). Thus, multiple lines of evidence suggest that the primate BG invigorates limb movements according to environmental and energetic constraints (Turner & Desmurget, 2010).

In this study, we explore the possibility that the importance of primate BG in limb vigor control extends to speech production. We recorded macroelectrode local field potentials and microelectrode single neuron activity from two key nodes in the BG, the subthalamic nucleus (STN) and the globus pallidus (GP), while patients undergoing deep brain stimulation (DBS) surgeries completed a speech-in-noise task. Using two measures of neural activity, local field potential broadband gamma activity (LFP-BGA) and single-unit firing rate (SU-FR), we aimed to answer two primary questions: 1) How do GP and STN differentially encode speech production? And 2) Do GP and STN encode speech-in-noise (‘Lombard’) modulations or any of its computational building blocks, such as ambient noise levels? The first question is motivated by the scarcity of high spatiotemporal resolution recordings from the BG during speech. STN activity has been described in several studies of speech and speech-like tasks (Chrabaszcz et al., 2021; Hebb et al., 2012; S. F. Johari et al., 2020; Tankus et al., 2021; Tankus & Fried, 2019; Watson & Montgomery, 2006), while GP speech modulation has not been described in the literature. Our cohort consisted of patient-participants undergoing either STN-DBS or GPi-DBS implantation, permitting comparison across targets. Our second aim was to characterize BG encoding of variables related to the Lombard effect, like speech intensity or ambient noise levels. Our main hypothesis was that populations in GP and STN correlate with speech intensity. Brücke et al. (2012) reported monotonic increases in the LFP-BGA in human GP with increases in arm movement amplitude (see also Tan et al. (2015) for similar results in finger force in the beta band). We hypothesized that these findings would generalize to speech, with soft speech evoking a small increase in LFP-BGA, medium speech evoking a modest increase in LFP-BGA, and loud speech evoking the largest increase in LFP-BGA.

We developed a permutation-based statistical analysis to account for non-stationarities observed in intraoperative recordings. We found that LFP-BGA and SU-FR were robustly modulated in both GP and STN in a sentence-repetition task. Task-related LFP-BGA increases in STN began immediately after stimulus offset in preparation for speech, while increases in LFP-BGA in GP occurred later and were more coincident with speech onset. All participants reliably increased voice intensity in response to background noise, but we did not find strong encoding of voice intensity in either nucleus—such correlations were limited to a small number of GP LFP-BGA recording sites and they occurred in the mid-sentence time window. Instead, we found that both LFP-BGA and SU-FR in both structures more robustly tracked ambient noise levels. From SU-FR population activity, we were able to decode the presence of background noise with ∼75% accuracy at any point in time over the course of the trial. Neural populations encoding ambient-noise and those encoding speech were independent; we found no relationship in LFP-BGA between the likelihoods of a recording site exhibiting speech modulation and it exhibiting ambient noise modulation. At the single neuron level, the populations were anticorrelated; speech-modulated neurons were less likely to be modulated by ambient noise compared with what was seen in the general population. Our results address important gaps in our understanding of basal ganglia contribution to speech production and neural coding of the Lombard effect. BG ambient noise level encoding is concordant with the idea that primate BG invigorate motor action, including speech, by setting an ‘urgency’ signal that then influences premotor and primary motor cortices’ selection and implementation of a motor plan (Thura & Cisek, 2017).

## Results

### Participants spoke louder in the presence of ambient noise in an intrasurgical sentence repetition task

We recorded basal ganglia (STN or GP) local field potentials and single-unit activity in a sentence repetition task in the operating room with 23 patient-participants (17 men, 6 women, mean age 63, age range [16-82]) undergoing deep brain stimulation implantation surgery (full demographic and disease characteristics are provided in Supplementary Table 1). The task, called the ‘Lombard’ task herein, asked participants to repeat aloud one of ten sentences (see Methods) on each trial. Trials consisted of three epochs: a baseline (BSLN) period in which participants rested, a STIMULUS epoch in which participants heard and read a sentence, and a SPEECH epoch in which participants repeated the sentence (Figure 1). Participants were asked to begin speaking after the text on the screen turned green. We defined the period after the go cue and before the SPEECH epoch as the reaction time (RT)/preparatory period. Participants produced sentences in one of two experimental conditions. In the NOISE condition (see Methods), 70 dB multitalker babble was played over loudspeakers with the intention of eliciting the Lombard effect. In the QUIET condition, no experimental background noise was presented. NOISE and QUIET conditions were presented in blocks of 10 trials. Up to eight blocks (four NOISE and four QUIET) were presented in each experimental run if the participants were able. Critically, the background multitalker babble was played continuously from the beginning of the first trial in the NOISE block to the end of the last trial (Fig 1A), resulting in two types of BSLN trials (BSLN-QUIET and BSLN-NOISE). The blocked design was intended to minimize the participants’ awareness of the experimental manipulation of the background noise of the task, and to mimic the timescale (∼minutes) of ambient noise changes in everyday environments.

**Figure 1:**
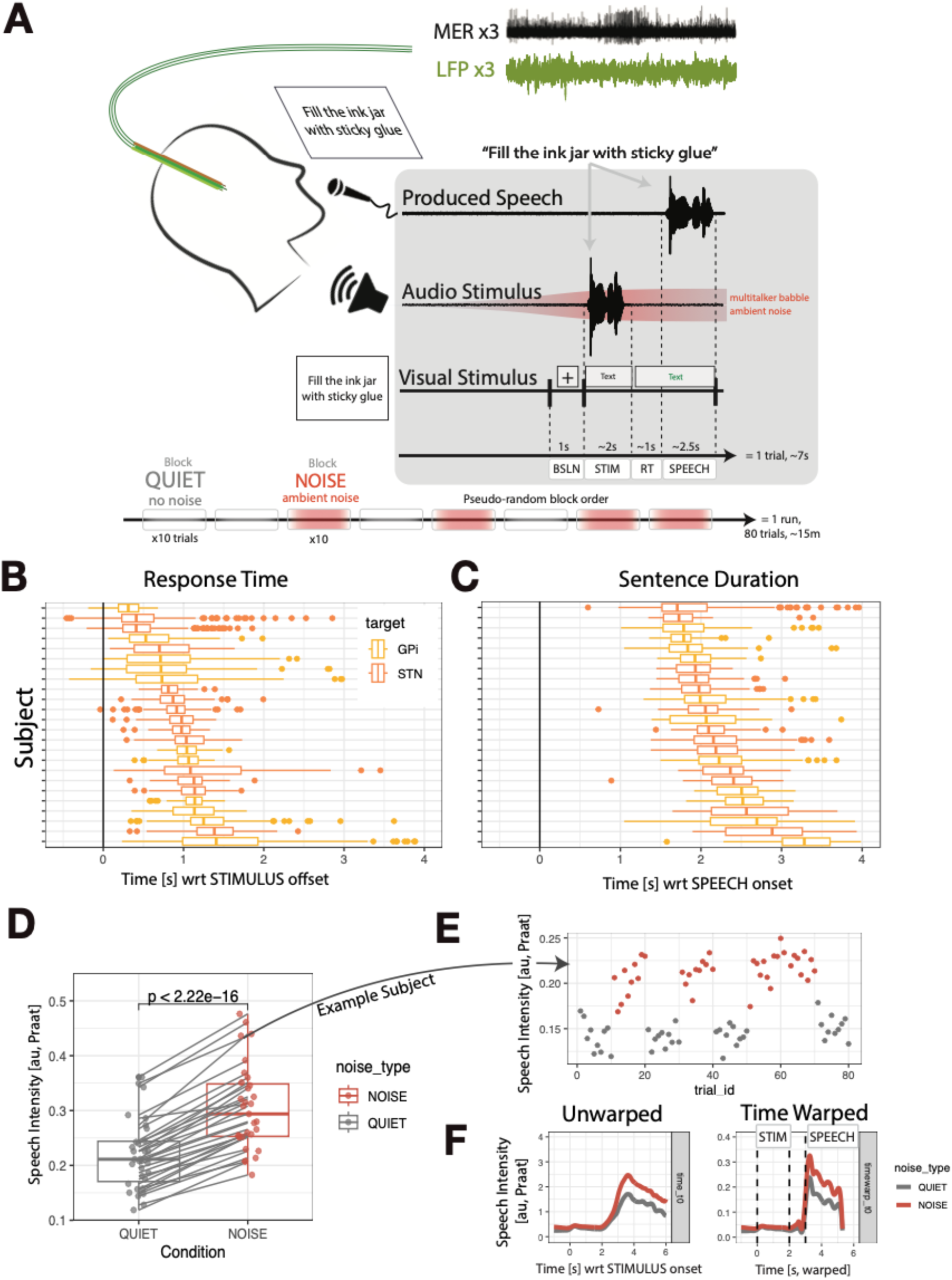
Ambient noise elicits an increase in speech volume during intraoperative sentence repetition task. (A) Experimental design of speech-in-noise sentence production. Patients in the operating room repeated sentences aloud in response to simultaneous auditory and visual stimuli (“Audio Stimulus” and “Visual Stimulus”). Every trial was divided into four epochs: baseline BSLN, stimulus STIM, response time RT, and speech SPEECH. In half of the trials, multitalker babble was played over loudspeakers (NOISE condition) with the intent of eliciting the Lombard effect. The other trials were QUIET. NOISE and QUIET trials were administered in blocks of 10 trials (bottom) in order to mimic everyday timescale of ambient noise changes. (B, C) Reaction time and utterance duration across the cohort. (B) Reaction time was defined as the duration of the PREP epoch. Subjects are sorted by median reaction time (left) and median sentence duration (right). Bars are color-coded by STN-DBS subjects (orange) and GPi-DBS subjects (yellow). Box-and-whisker plots indicate median and lower/upper quartiles. Outliers are plotted as dots. (C) Same as (B) for sentence duration. (D) Voice intensity in the QUIET vs NOISE condition. Each point in a condition represents the average intensity across trials for one participant. Note the positive average slope from QUIET to NOISE condition, indicating a consistent and pronounced speech-in-noise (‘Lombard’) effect. (E) Example run from one participant showing the modulation of voice intensity by ambient noise levels. Each point represents average voice intensity during the voiced periods of a given trial. Red dots were produced in the NOISE condition, and grey dots were produced in the QUIET condition. Note the higher intensity values during the NOISE blocks relative to the QUIET blocks. (F) Time warping to align task events across trials and participants. Left, grand average voice intensity curves (as estimated by PRAAT) in the QUIET and NOISE conditions time-locked to the beginning of each trial (STIMULUS onset). Right, grand average voice intensity curves after linear time-warping to key events: STIMULUS onset (t=0), STIMULUS offset (t=2), SPEECH onset (t=3), and SPEECH offset (t=5.3). Note the sharper rise at speech onset for the right plot relative to the left, indicating proper alignment of the behavioral events.

Participants completed 105 ± 56 [18-237] (mean ± sd [range]) trials across 1-3 runs of the task. As the experiment was intended to elicit the Lombard effect, we first analysed whether participants spoke with higher vocal intensity (radiated acoustic sound pressure measured in dB SPL, henceforth ‘speech loudness’) in the NOISE versus QUIET condition (Fig 1D). At the cohort level, participants spoke louder in the NOISE condition (paired t-test, p=2.22e-16). All (23/23) participants showed the effect (unpaired t-test within participant, alpha=0.05 threshold; Fig 1E).

Although an increase in voice intensity is the classic manifestation of the Lombard effect, previous studies have also reported other concomitant changes in speech in response to an increase in ambient noise levels (see Introduction; (Brumm & Zollinger, 2011)), including increased fundamental frequency (Stowe & Golob, 2013), vowel duration, and the size of a speaker’s vowel space (Elie et al., 2024). We compared the NOISE and QUIET conditions using these measures at the participant level. In the NOISE condition we observed increased fundamental frequency (Fig 1—S1; paired t-test p=4.2e-9); increased vowel duration (Fig 1—S1; paired t-test p=2.6e-8); and expanded vowel space as measured by the formant centralization ratio (Sapir, 2014) (Fig 1—S1; paired t-test p<1.4e-4). Lastly, a separate, non-acoustic channel reinforced evidence of speech-in-noise changes in the NOISE condition; respiratory excursion, as measured by a force transduction belt over the rib cage, was larger in the NOISE vs QUIET condition (Fig 1—S2). We thus elicited a constellation of speech-in-noise behavioral changes by manipulating ambient noise levels. Because it was the most robust measure of NOISE vs QUIET, voice intensity is used as the primary index of the Lombard effect in all subsequent neural analyses.

Reaction time and speech production duration were highly variable (Fig 1--S3), resulting in trial durations as short as 3.5 s and some as long as 6 s (Fig 1B-C). To align data across all trials and participants, we used event-based linear time warping according to mean event times across the cohort (Fig 1F). The STIMULUS period was 1.97 ± 0.21 s (mean ± sd), RT was 0.97 ± 0.26 s, and SPEECH was 2.3 ± 0.38 s. For across-participant analyses, we rounded these times and linearly warped all time-series behavioral and neural signals such that, with respect to STIMULUS onset, every trial aligned with the following landmarks: STIMULUS onset at time 0, STIMULUS offset at 2 seconds, SPEECH onset at 3 seconds, and SPEECH offset at 5.3 seconds (see (Dichter et al., 2018; Kobak et al., 2016) for other examples of time-warping in neurophysiological data). When applied to the time traces of produced acoustic intensity, the time warping approach effectively aligned signals across trials and subjects, sharpening the increase of acoustic intensity at SPEECH onset and sharpening the decrease at SPEECH offset (Fig 1F).

To test if disease severity influenced behavior in the Lombard task, we used the subset of the cohort diagnosed with Parkinson’s disease to analyze whether Lombard effect size (mean voice intensity in the NOISE minus the QUIET condition) was correlated with preoperative Movement Disorder Society-Unified Parkinson’s Disease Rating Scale (MDS-UPDRS) part III scores (Fig 1--S4). We found no relationship between preoperative off-medicine MDS-UPDRS part III scores and Lombard effect size (Spearman correlation, p=0.73), nor between the dopaminergic medicine MDS-UPDRS part III change (off-medicine minus on-medicine) and Lombard effect size as measured by speech loudness (Spearman correlation, p=0.42).

Participants with STN and GP targets exhibited similar reaction times and utterance durations (Fig 1—S5; Wilcoxon rank-sum, reaction time p=0.27, utterance duration p=1.0). This similarity rules out gross behavioral differences as major drivers of differences in STN and GP neural activity examined in subsequent analyses.

In summary, our experimental design robustly elicited louder speech in the NOISE relative to the QUIET condition.

### STN and GP are modulated by speech production; STN activates prior to GP at the LFP level

In each recording session, we recorded neural activity from two to three left-hemisphere clinical microelectrode tracts targeting either the STN (n=12 participants) or GPi (n=11 participants; Fig 2A) during the microelectrode mapping stage of DBS surgery (see Methods). We used recording probes (Sonus, AlphaOmega) that had 1) standard clinical microelectrodes from which we recorded single-unit (SU) activity and 2) macroelectrode rings 3 mm superficial to the tip from which we recorded LFPs (Fig 2A, right). Due to their spacing, the macroelectrode LFP recordings and the microelectrode SU recordings were *not* from the same locations on any given run of the task. For STN cases, it was common to record LFPs from outside the atlas-defined STN border (Fig 2F, all dots) since they were 3 mm superficial to the clinical microelectrode target. Similarly, some LFP recording sites from GPi-DBS cases were localized to the *external* segment of the GP.

**Figure 2:**
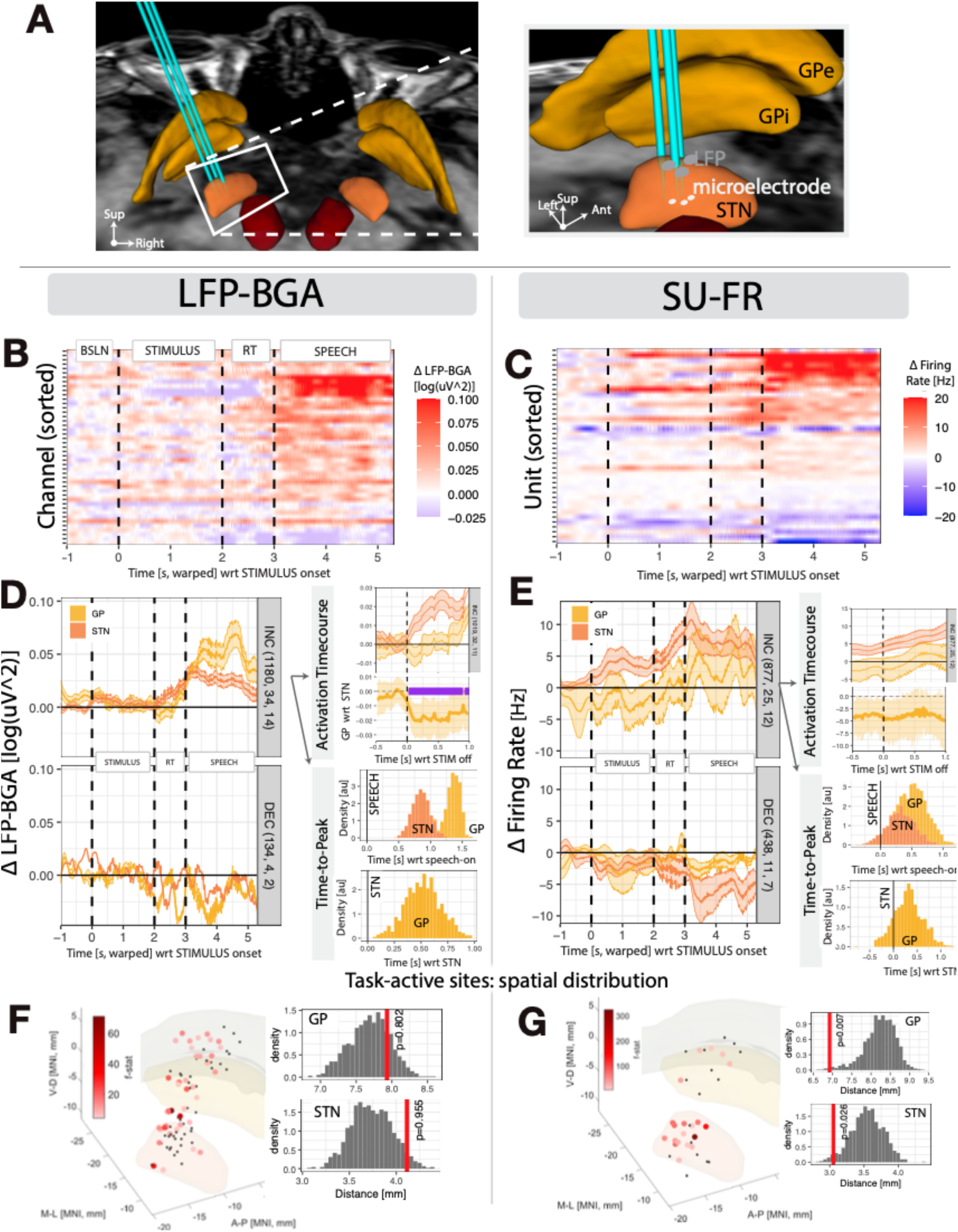
STN and GP activity are modulated by speech production. (A) Example native-space reconstruction of surgical stimulation tracks and recording sites in one patient-participant receiving STN-DBS. (Left) Posterior view of the tracks and nuclei of interest: internal and external segments of the GP (light orange) and the STN (dark orange). Red nucleus (RN) is also visualized as a landmark. Clinical microelectrodes along with research local field potential rings were inserted along 3 tracks (teal). (Right) Medial view zoomed into the STN. LFPs were recorded 3 mm superficial to the microelectrode tips along each track. In this example subject, the microelectrodes recorded from mid-to-ventral STN (white circles), while the macroelectrodes recorded from dorsolateral STN (grey circles). (B,D,F) Local field potential broadband gamma activity (LFP-BGA). (C,E,G) Same analyses as B,D, and F using the single-unit firing rate (SU-FR) recordings. (B) Raster plot of activity for all task-active channels. Pixel color indicates LFP-BGA log-power averaged across trials at a given channel and time bin. Channels are sorted by sign and magnitude of activation from top to bottom. Note the activation (red) across many channels starting at SPEECH onset (t=3) and continuing for the duration of the production. (C) Same as (B) for SU-FR. Note the positive (red, top) and negative (blue, bottom) changes in firing rate. (D) Time course of activation for STN (orange) and GP (yellow) channels, split by channels with increasing (INC) and decreasing (DEC) deflections in the SPEECH window with respect to baseline. Right, “Speech Onset”: bootstrapped GP-versus-STN (mean ± 95% CI, across channels) response profile of INC-type recording sites time-locked to speech onset, with significance (alpha < 0.05) indicated in purple. STN activated just after STIMULUS-offset, while GP dipped and slowly ramped up to speech. Right, “Peak of activation”: time of activation peak in GP and STN (and GP vs STN) time-locked to speech onset bootstrapped across channels. (E) Same as (D) for SU-FR. (F) Spatial distribution of task-active recordings sites in MNI space. DISTAL atlas STN, GPi, and GPe are plotted from foreground to background. Refer to panel A, right, for orientation and other landmarks. Circles represent recording sites. Red circles were significantly task-activated, with degree of activation (measured by f-statistic) indicated by the intensity of red. Right: spatial clustering test of task-activated channels with respect to all channels for GP and STN. Red lines indicate *actual* mean distance between task-active channels (averaged over all pairwise connections). Null distributions calculated with shuffled active/inactive labels. (G) Same as (F) for SU-FR. Data quantities are indicated as (# trials, # channels/units, # participants) in each panel. Vertical lines in time-series plots represent: STIMULUS onset, STIMULUS offset, and SPEECH onset.

After spike sorting, artifact rejection, and data exclusion (see Methods), the final cohort consisted of 23 participants, with 100 macroelectrode LFP channels and 57 single units (see Table S1 for a tabulation of channel and single unit counts). We categorized units as stationary single-units (n=22), non-stationary single-units (n=29), stationary multi-units (n=13), or non-stationary multi-units (n=31) based on published methods (Lipski et al., 2018; Pouzat et al., 2002; Rutishauser et al., 2006a), including rigorous manual unit-by-unit inspection (see Methods). We did not initially exclude units on the basis of this categorization, but later used it in assessing robustness of SU analyses. Similar qualities of units were recorded across GP and STN (Fig 2--S1).

Many LFP recording sites were near the border between the internal segment and external segment of the GP. Because intracranial recordings during speech have never been reported from either segment of the GP to our knowledge, we opted to combine all pallidal recordings for a tractable analysis in comparing the GP with STN. The final electrophysiology dataset used in statistical analysis, after rejecting runs in each modality separately that didn’t have at least 10 trials, consisted of 51 GP and 49 STN LFP recording sites, and 23 GP and 35 STN single units.

We observed task-related modulation across the LFP and single unit firing rates in both GP and STN (Fig 3B, C). Beta desynchronization is the most salient LFP modulation in task-related basal ganglia recordings. We verified that the LFPs exhibited strong beta-band desynchronization and exhibited the expected 1/f power spectral density profile (Donoghue et al., 2020; Kramer & Chu, 2024; Pritchard, 1992) (Figure 2--S1). We subsequently focused on high-frequency activity (> 40 Hz) because previous studies found that gamma activity scales with limb movement velocity (Brücke et al., 2008; Fischer et al., 2017; Jenkinson et al., 2013; Joundi et al., 2012; Lofredi et al., 2018) and because we have previously observed speech-related activity in STN in that frequency band (Chrabaszcz et al., 2019). The bandwidth of prominent speech-related activation varied across electrodes, with some modulations confined to the 60-120 Hz and others extending up to 250 Hz. We thus decided to extract broadband gamma activity (60-250 Hz) from the LFP with multitaper DPSS spectral decomposition (Prerau et al., 2017). We use this one-dimensional signal to investigate coding of speech and ambient noise levels in all subsequent LFP analyses.

**Figure 3:**
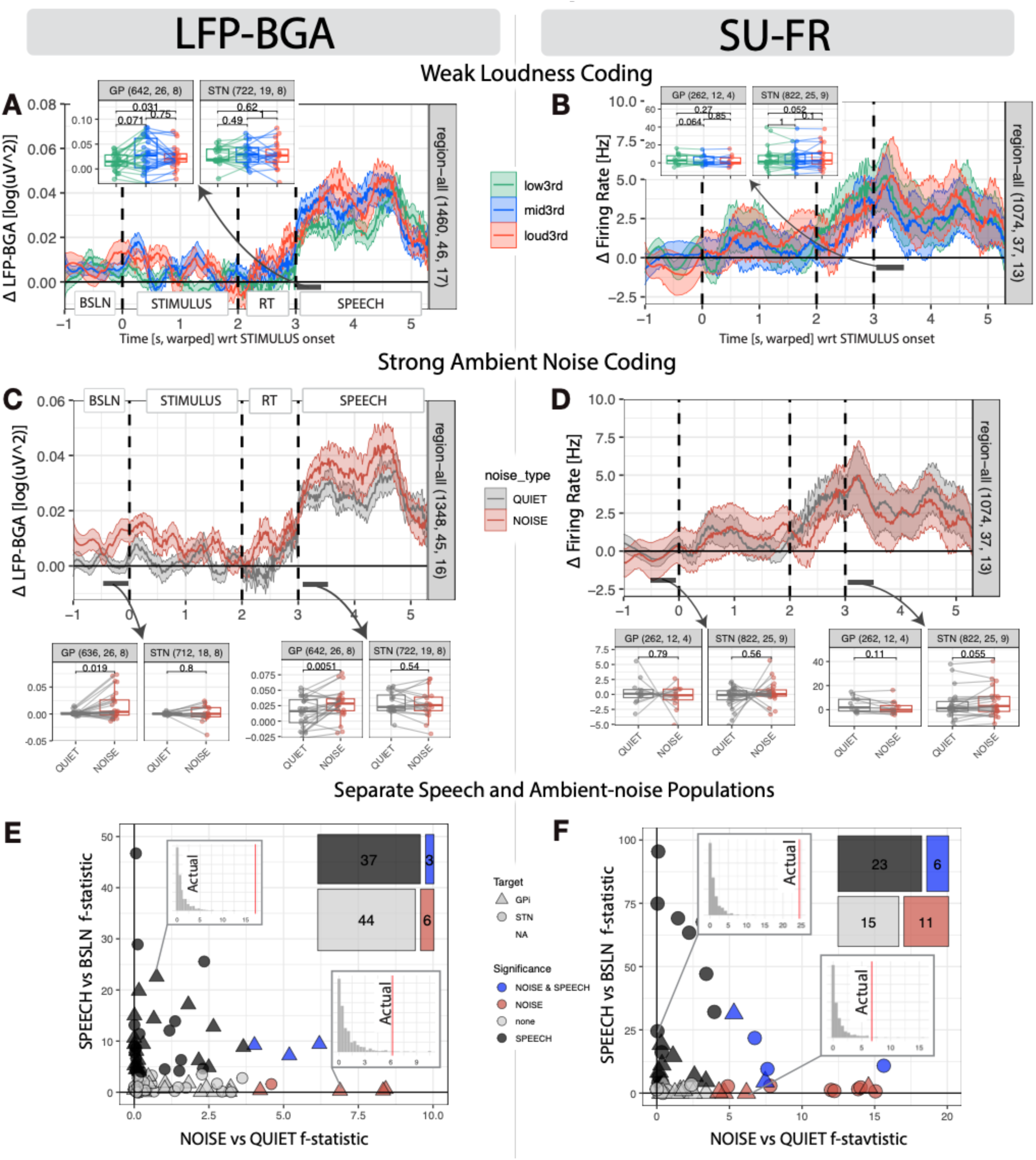
Speech-modulated and ambient-noise-modulated populations were distinct. Panels are organized by modality: LFP-BGA on the left, and the same analyses using SU-FR on the right. (A, B) LFP-BGA and SU-FR activation split by produced-speech loudness levels determined within-participant. Traces represent mean activation within the quietest, middle, or loudest trials within each participant. Shaded errors represent S.E.M. Few consistent differences were observed across conditions. Inset plots are the same data split into STN and GP and collapsed into a single point by averaging the first 500 ms of SPEECH. Each point represents the average for a single run of the task for a participant; box-and-whisker plots summarize the data with median, upper and lower quartiles. Pairwise Wilcoxon-signed rank tests revealed no notable trends in LFP-BGA (low3rd vs mid3rd p=0.071, mid3rd vs loud3rd p=0.750, low3rd vs loud3rd p=0.031 for GP; low3rd vs mid3rd p=0.490, mid3rd vs loud3rd p=1.00, low3rd vs loud3rd p=0.62 for STN) or in SU-FR (low3rd vs mid3rd p=0.064, mid3rd vs loud3rd p=0.85, low3rd vs loud3rd p=0.27 for GP; low3rd vs mid3rd p=1, mid3rd vs loud3rd p=0.1, low3rd vs loud3rd p=0.052 for STN). (C, D) Same as (A, B) but split by ambient NOISE (red) and QUIET (grey) condition rather than produced loudness. (C) LFP-BGA was consistently higher across in the NOISE condition relative to the QUIET condition. Inset plots are split by region (GP and STN) and averaged in the BSLN window [-0.5, 0] s (left) or the first 0.5 s of SPEECH [3, 3.5] s (right). In GP, but not STN, NOISE was greater in both tested time windows (Wilcoxon signed-rank in GP, BSLN p=0.019 and SPEECH-beg p=0.0051). (D) The ambient noise condition did not have an effect on SU-FR; no regions across any time windows were significant (Wilcoxon signed-rank). (E, F) Single-channel linear modeling with permutation-based significance testing. Neural data were modeled with a 2x2 design: SPEECH (average [0, 0.5] s with respect to speech onset) vs BSLN (average [-0.5, 0] s with respect to STIMULUS onset) and NOISE vs QUIET. Each point in the scatter plot represents a channel (LFP-BGA) or single neuron (SU-FR). Y-axis indicates each channel’s f-statistic for the SPEECH vs BSLN model coefficient. X-axis indicates each channel’s f-statistic for the NOISE vs QUIET model coefficient. Larger values indicate strength of modulation in SPEECH relative to BSLN and NOISE relative to QUIET. Insets visualize example channels: actual f-statistic (red vertical line) relative to the histogram of that channel’s null distribution (light grey). Top right, contingency tables: counts of each category of modulation. Data quantities are indicated as (# trials, # channels/units, # participants) in each panel.

Grand-average LFP-BGA increased relative to baseline during the speech preparatory window and was sustained through the middle and end of the utterance (Fig 2D). Neither STN or GP LFP-BGA activated in response to the stimulus. Just after STIMULUS offset—in the preparatory period prior to speech production—STN activated swiftly while GP more slowly ramped up to speech onset (Fig 2D, top right). In a direct comparison (sliding window bootstrap across-channels, two-sided alpha<0.05), GP and STN LFP-BGA diverged immediately at STIMULUS offset for approximately 1 second thereafter. We also quantified the time of peak activation in LFP-BGA (Fig 2D, bottom right); both STN (95% CI [0.58, 1.13] s, mean 0.85 s) and GP (95% CI [1.18, 1.54] s, mean 1.3 s) peaked after speech onset. GP peaked after STN (95% CI [0.13, 0.82] s, mean 0.52 s).

SU firing rates were also task-modulated (Fig 2C). In contrast to LFP-BGA deflections, which were positive, SU-FR changes were both positive and negative relative to baseline, which we refer to as “increasing-type” (INC) and “decreasing-type” (DEC), respectively, following the categorization of previous work (Lipski et al., 2018; Vissani et al., 2025). Increasing and decreasing types were assigned to each unit according to the sign of the greatest absolute coefficient of sliding-window bootstrapped mean differences in the -2 s to 2 s window around SPEECH onset (see Methods). There were 26/58 INC SUs and 12/58 DEC SUs, leaving 20/58 SUs that were not significantly task-modulated. Other studies have also characterized mixed-type SUs (Lipski et al., 2018) but, for the sake of tractability, we will adhere to the binary (INC/DEC) categorization and focus on the INC type SU for appropriate comparisons with LFP-BGA increases. Unlike LFP-BGA responses, SU-FR displayed a clear STIMULUS epoch modulation, with SUs decreasing or increasing firing rate coincident with stimulus presentation (Fig 2C). STN SU-FR time-to-peak was coincident with SPEECH onset (bootstrap across units, 95% CI [-0.11, 0.77] s, mean 0.33 s). GP SU-FR time-to-peak was after SPEECH onset (Fig 2E, right; bootstrap across units, 95% CI [0.29, 1.01] s, mean 0.64 s). In a direct comparison, GP SU-FR was not significantly different from STN SU-FR time-to-peak when time-locking to SPEECH onset (Fig 2E, right; bootstrap across units, 95% CI [-0.25, 0.89] s, mean 0.31 s).

LFP-BGA activations were not spatially concentrated in any subregion of either STN or GP relative to the overall distribution of sampled sites (Fig 2F; spatial aggregation permutation test, p=0.995 and p=0.802 for STN and GP respectively). SU-FR task-active sites, on the other hand, were spatially concentrated relative to the overall distribution of sampled sites (Fig 2G; spatial aggregation permutation test, STN p=0.026 and GP p=0.007). In STN, task-active sites (centroid MNI (x,y,z) =(-14.2, -14.3, -7.5)) trended more laterally and dorsally than would be expected by chance (Fig 2G; all sites centroid (-13.2, -14.5, -8.3)). In GP, task-active sites (centroid MNI (x,y,z) =(-16.3, -12.2, -5.7)) trended more laterally than would be expected by chance (Fig 2G; all sites centroid (-16.6, -11.4, - 5.8)).

### Distinct Speech- and ambient-noise-modulated neural populations

We hypothesized that neural populations in GP and STN would encode voice intensity. In particular, we expected louder trials to be associated with greater activation of GP and STN (as motivated in the Introduction). To test this, we inspected how LFP-BGA changed with different speech intensities (Fig 3A,B). Trials were sorted according to their voice intensity and divided into tertiles: loud, middle, and soft. We tested the intensity-BGA relationship in the first 500 ms of the SPEECH window, keeping in line with previous research showing GPi encoding of movement intensity starting at movement onset and persisting for hundreds of ms thereafter (Brücke et al., 2012). In both LFP-BGA and SU-FR, there were no robust differences in neural activity between loud and soft trials (Fig 3A,B insets, Wilcoxon signed-rank p>0.01). In LFP-BGA, we observed a medium > loud > soft trend in GP (Fig 3A inset). We identified a single participant with clear, consistent differences between loudness tertiles in LFP-BGA in the hypothesized direction.

Finding only weak evidence supporting our primary hypothesis, we explored the possibility that speech loudness is coded in other time windows or in specific regions, using the same approach as above (Fig 3--S2). We found that: 1) GP, and not STN, showed evidence of loudness encoding and 2) these effects were most prominent in the mid-sentence time window (Wilcoxon signed-rank, low3rd vs loud3rd in the 3.5 - 4 sec time window, GP p=0.00053 vs STN p=1.0). This effect held even when potential outliers were removed (GP p=0.0072 vs STN p=1.0).

Our previous analyses split neural activity by behavior. Interestingly, when we split trials by environmental condition, we noted prominent group-level NOISE vs QUIET differences in LFP-BGA (Fig 3C). LFP-BGA was consistently greater in the NOISE condition relative to QUIET condition throughout the whole trial, including the baseline, suggesting that LFP-BGA was indexing an environmental variable (high ambient noise levels vs low ambient noise levels) rather than behavior (speaking loud vs soft). Statistical analysis confirmed that 1) there was indeed a NOISE vs QUIET effect and 2) the effect was specific to the GP (Fig 3C bottom left, Wilcoxon signed-rank p=0.0051 in GP, p = 0.65 in STN). This effect was also independent of task epoch; the effect was present whether we looked in the BSLN, STIMULUS, or SPEECH window (Fig 3C, bottom left and bottom right, SPEECH window Wilcoxon signed-rank p=0.0051 in GP). As a control, we verified that this NOISE vs QUIET effect was abolished when we shuffled NOISE and QUIET trial labels (Fig 3--S3). Thus, GP LFP-BGA power tracked ambient noise levels in addition to being speech-modulated.

At the group average level, there was no consistent SU-FR modulation for the NOISE condition in either the STN or GPi (Fig 3D; Wilcoxon signed-rank p>0.05 for all comparisons). However, the channel-averaged group analyses didn’t address a) whether there may be both negative and positive NOISE modulations at the single-channel or single-unit level and b) whether SPEECH-modulated and NOISE-modulated neural populations were interdependent or independent populations. We thus designed a permutation-based statistical approach that operated on each channel individually, leveraging inter-trial variability and trial shuffling to determine statistical significance (see Methods). The approach consisted of two phases: 1) modeling neural activity as a function of task epoch (SPEECH vs BSLN) and ambient noise levels (NOISE vs QUIET) with a generalized linear model, followed by 2) generation of null distributions for each predictor via shuffling. We shuffled model *residuals* rather than model *predictors*, which is a particularly conservative approach to ensure that the null distributions are truly comparable to the actual values. Significance levels for SPEECH (relative to BSLN) and NOISE (relative to QUIET) for each neuron (SU-FR) or channel (LFP-BGA) were defined by how unlikely the actual data predictor coefficients were relative to the null distribution of coefficients.

In LFP-BGA, we identified 44% (40/90) channels activated during SPEECH relative to BSLN (Fig 3E). 10% (9/90) showed a difference for NOISE relative to QUIET trials, whose actual f-statistics were beyond the null distribution of f-statistics (Fig 3E inset). We hypothesized that the NOISE-modulated populations would be a subset of the SPEECH-modulated populations, which would drive a linear positive correlation between the two f-statistics at the population level. We instead observed a largely independent relationship between the SPEECH and NOISE f-statistics (Fig 3E scatter). We quantified this relationship with a contingency table by tabulating significant channels (Fig 3E top right inset). We found no significant dependence between the likelihood any given channel was SPEECH-modulated and the likelihood it was NOISE-modulated, suggesting these two populations are independent (Barnard’s test, p=0.37).

The same analyses with SU-FR revealed that 41% (23/55) of neurons were SPEECH modulated (consistent with the LFP-BGA results). However, a higher proportion of neurons showed a difference for NOISE relative to QUIET trials: 31% (17/55) compared to 10% in LFP-BGA. SU-FR traces confirmed these results and also revealed the variability in the sign of the effect (Fig 3--S1), with some neurons characterized by a NOISE > QUIET offset and others characterized by a QUIET > NOISE offset. Six neurons were both SPEECH and NOISE modulated, leaving eleven that were NOISE-only modulated. SU-FR NOISE modulations were significantly *more* likely in neurons that were not SPEECH modulated (Barnard’s test, p=0.048). That is, the presence of SPEECH modulation in a given neuron’s firing rate made it less likely that the neuron was NOISE modulated.

### Population analysis demixes and decodes speech-related and ambient-noise-related modulations

Prior analyses used group-level channel-averaged statistics (Fig 3C,D) and subsequently channel-wise single-trial models (Fig 3E,F) to investigate speech and ambient noise modulation in BG. The group-level channel-averaged approach (Fig 3C-D) was time-resolved but is limited in statistical power and cannot inspect relationships between speech and noise modulations. On the other hand, the trial-level approach (Fig 3E-F), gives channel-wise statistics—conferring speech and noise modulation population analyses—but is not time-resolved and fails to leverage all data in a single model. We thus turned to an approach that can balance these factors: demixing PCA (dPCA) (Kobak et al., 2016). dPCA is a well-established technique in experimental cognitive neurophysiology to dissect neural encoding of experimental conditions and behavior. dPCA is like classic principal components analysis (PCA) in that it decomposes a data matrix into a set of components, but diverges starkly from PCA in that dPCA components are not necessarily orthogonal, and dPCA has different ‘encoder’ and ‘decoder’ axes. dPCA also doesn’t attempt to reconstruct the full data matrix. Rather, the dPCA optimization loss function operates over *marginalizations* of the data according to experimental conditions.

We expected dPCA to find significant components that reflect ambient noise tracking. One advantage of dPCA is that it can account for variability in the polarity of responses by finding a neural subspace that has negative weights for some units and positive weights for others. This contrasts with, for example, a linear mixed effects model whose fixed-effect sensitivity requires all within-participant coefficients to be consistently positive or consistently negative. For this reason, we hypothesized that dPCA would more effectively reveal single-unit coding of ambient noise compared to previous analyses (Fig 3).

For each modality (LFP-BGA and SU-FR), we aggregated the dataset across all regions and participants into a pseudo-population in line with previous studies (Fetterhoff et al., 2024). We configured dPCA to decompose the data into axes concerning two aspects of the data: time-resolved neural activation independent of condition (ie, stimulus- or speech-related activity), and ambient noise coding according to task condition (Fig 4).

**Figure 4:**
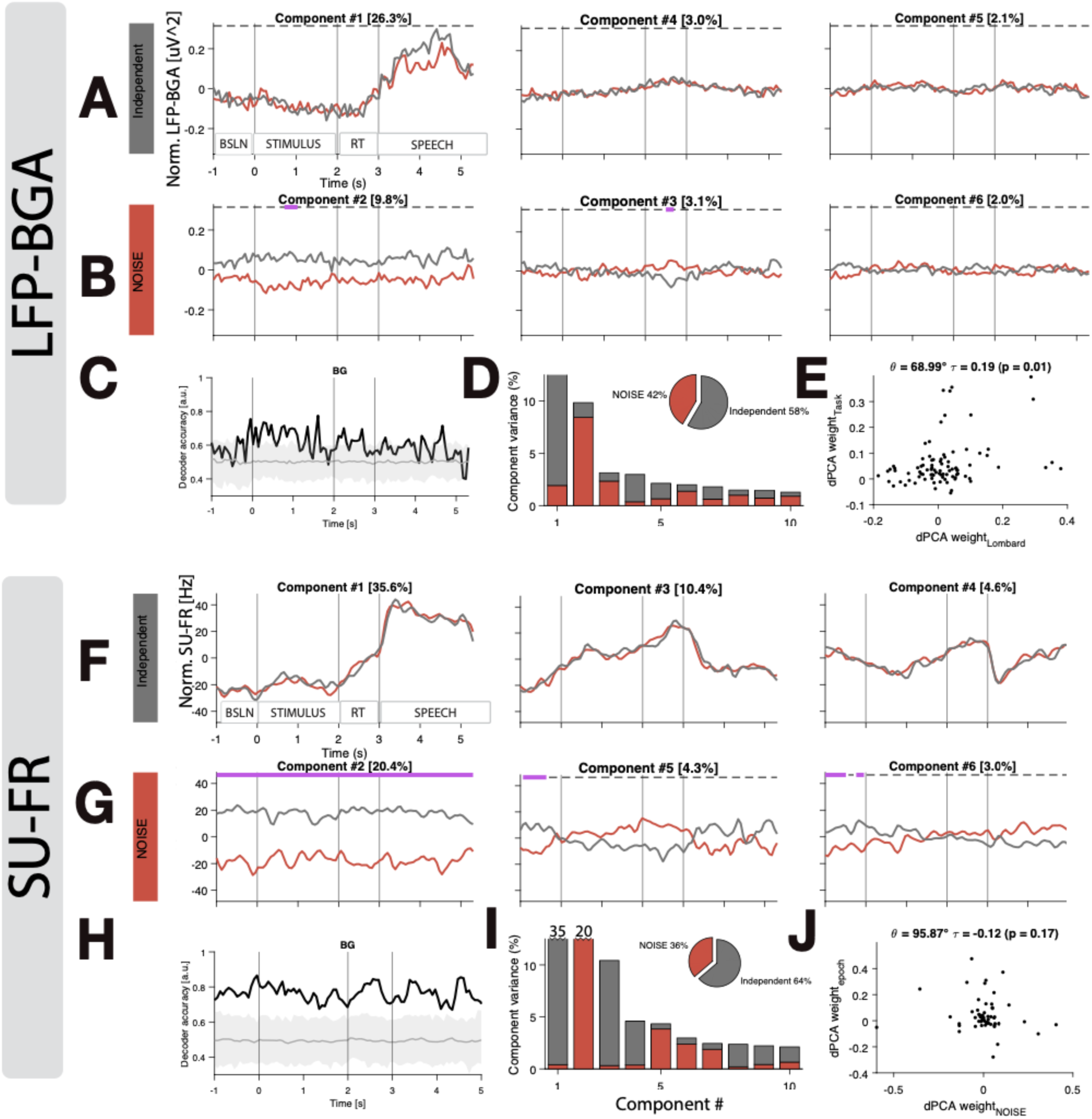
dPCA reveals strong population encoding of ambient noise levels at the single neuron level. dPCA population-level approach to identifying task-related neural coding. (A, F) dPCA condition-independent components, sorted left to right by percentage explained variance. Lines represent population average traces of NOISE (red) and QUIET (grey) after projection onto the component. (B, G) NOISE-related components, as in (A, F). Lines at the top of each panel indicate whether the component was significant in a specific time window: dashed indicates insignificant time windows while solid purple indicates significant time windows. (C, G) Accuracy over time (black) of linear decoder classifying NOISE vs QUIET in data projected onto the NOISE component. Shaded grey region indicates 95% confidence bounds from a shuffle test—accuracy above this was considered statistically significant. (D, H) Each dPCA component (x-axis; sorted by explained variance), its percentage of explained variance (y-axis), and how each marginalization contributed to the component (red/grey proportion within each bar). Inset pie chart aggregates percentage of explained variance across all components. (E, I) Relationship between encoding weights for the first condition-independent component (y-axis) and the first NOISE-related component (x-axis). Points represent LFP-BGA recording sites (E) or neurons (J). The angle between the encoding vectors reveals how SPEECH and NOISE were encoded across the population (angle = 67°, Kendall correlation p=0.01 for LFP-BGA; angle = 96°, Kendall correlation p=0.17 for LFP-BGA). In plots with time on the x-axis, the three vertical lines represent: STIMULUS onset, STIMULUS offset, and SPEECH onset

The component that explained the largest portion of variance in the data was the speech-related component (“condition-independent” in dPCA terminology) in both LFP-BGA (26.3%) and SU-FR (35.6%; Fig 4A,E left). dPCA found a similar time course of activation in the task for LFP-BGA as in previous group analyses (Fig 3C), characterized by the following dynamics relative to the baseline epoch: 1) weak deactivation in the STIMULUS epoch, 2) activation in the RT/PREP epoch starting ∼500 ms prior to SPEECH onset, 3) peak activation in the middle of SPEECH production, and 4) a steady ramp down of activity towards the end of SPEECH (Fig 4A). For LFP-BGA, the second and third condition-independent components accounted for little variance (3.0% and 2.1%), suggesting that many recording sites exhibited similar activation profiles and were accounted for by the first component. SU-FR responses, in contrast, relied more on the second and third condition-independent time course to account for variability in responses (at 10.4% and 4.6% variance, respectively), reflecting the greater diversity of single-unit timecourses.

dPCA revealed stronger ambient noise coding in SU-FR than in LFP-BGA (Fig 4B, F). In LFP-BGA, although the magnitude difference between NOISE and QUIET components is evident when averaging across channels (Fig 4B red vs grey lines), we were not able to robustly decode the NOISE vs QUIET condition over the trial time course (Fig 4B purple, Fig 4C black). The first NOISE component was a stable offset that was largely independent of time in both modalities (which is not by methodological design (Kobak et al., 2016)). The absence of temporal dynamics reinforces the idea that neural populations are tracking ambient noise on the scale of tens of seconds or greater levels rather than moment-to-moment fluctuations. The latter would drive increases in the STIMULUS and SPEECH trial windows, and we did not find such transient changes. Note that the polarity of the axes is not meaningful in dPCA components; the QUIET condition being the higher of the two traces in the first NOISE-related component does *not* imply that SU-FR and LFP-BGA was higher across all recording sites in the QUIET condition. Rather, it simply means that SU-FR and LFP-BGA were *different* along the projected axes, with some recording sites being higher—and some lower—in the QUIET condition. We did not consider the second and third NOISE components meaningful as they comprise a small portion of variance (<5%; Fig 4B, F middle and right panels).

The NOISE-related components comprised a similar overall proportion of data variance in LFP-BGA and SU-FR (Fig D, I inset pie chart). It is worth noting, however, the distribution of that accounted variance across components in LFP-BGA versus SU-FR; the first NOISE-related component accounted for only 9.8% of overall variance for LFP-BGA (Fig 4B, D), while it accounted for 20.4% of variance for SU-FR (Fig 4G, I). That is, NOISE components accounted for a higher and more separable portion of variance in SU-FR compared to LFP-BGA.

Linear decoding from the data projected onto the NOISE axes reveals how strongly the condition is represented in the data. In LFP-BGA, despite decoding accuracy of NOISE vs QUIET staying >50% for nearly the whole timecourse of the trial, few individual timepoints reached the 95% significance threshold determined by shuffled data (Fig 4C). In SU-FR, decoding accuracy was higher and more consistent than for LFP-BGA, exceeding the significance threshold for the entire duration of the trial window (Fig 4G). Average decoding accuracy was 77% across the trial.

We inspected the relationship between SPEECH coding and NOISE coding by looking at the distribution of dPCA encoding weights in recording sites (LFP-BGA; Fig 4E) or neurons (SU-FR; Fig 4J). From prior analyses (Fig 3), we expected to find that separate populations of LFP and single-unit sites respond to speech, and another population tracks ambient noise levels. We used the first condition-independent component encoding vector, which we took as a proxy for SPEECH encoding, and the first NOISE component encoding vector. A linear correlation between values in these two vectors would indicate that SPEECH-related neurons tend to also be NOISE-related (and vice versa), while no correlation would indicate independence of the two encoding populations. In SU-FR, we observed no relationship between SPEECH and NOISE encoding (Fig 4J; Kendall rank correlation in line with (Kobak et al., 2016), p=0.17). In LFP-BGA, we observed a correlation between the two encoding vectors (Fig 4E; Kendall correlation, p=0.01), while noting the presence of apparent outliers in the distribution.

dPCA validated and extended previous analyses by providing a data-driven, non-parametric, population-level approach to studying both condition-independent and condition-dependent neural activation in the sentence repetition task. Most importantly, dPCA showed that ambient noise level can be robustly decoded from SU-FR and weakly decoded from LFP-BGA in GP and STN.

## Discussion

We investigated the neural underpinnings of speech production and the Lombard effect—often defined as an involuntary increase in voice intensity in response to background noise—in two key regions of the basal ganglia using rare single-unit and local field potential recordings from awake deep brain stimulation surgeries. Our intraoperative sentence-repetition Lombard task robustly elicited an increase in speech loudness, as well as changes in other measures of speech gain, in blocks of trials where multitalker babble background noise was presented to participants (Fig 1; Fig 1--S1&S2). With a stark scarcity of studies reporting direct time-resolved electrophysiological recordings of GP during speech production, we first aimed to coarsely characterize GP LFP-BGA and SU-FR timing with that of the STN. GP activity followed that of STN and peaks after speech onset (Fig 2), suggesting it does not contribute to speech initiation or sound sequencing.

Second, we aimed to understand how the Lombard effect is coded in basal ganglia. Inspired by studies of human and non-human primate coding of limb gain control (Anzak et al., 2012; Brücke et al., 2012; Tan et al., 2016), we hypothesized that neural activity in GP and STN would show Lombard coding in the first ∼500 ms of speech production, wherein LFP-BGA power monotonically increases as loudness increases. We found weak evidence of such coding, limited to GP LFP-BGA in the mid-sentence time window and only clearly present in ∼3 recording sites (Fig 3; Fig 3—S2). Instead, we found stronger evidence that GP encodes ambient noise level, the key environmental parameter driving the Lombard effect, in LFP-BGA (Fig 3) and SU-FR (Fig 3F, Fig 4). Our findings are highly concordant Thura and Cisek (2017), in which monkeys engaged in high-urgency and low-urgency blocks of a task, akin to our NOISE and QUIET blocks. GP neurons increased or decreased firing rate for the whole duration of urgency blocks, even in the baseline of the task.

These conclusions are bolstered by multiscale (single unit and local field potential) recordings and robust, permutation-based statistical modeling that accounted for neural non-stationarities.

### The lombard effect: voice intensity, vowel duration, vowel centralization, pitch, and respiration depth

Our behavioral findings reveal that the Lombard effect, as elicited during intraoperative sentence repetition, reflects a coordinated, cross-system adaptation rather than a unidimensional increase in vocal intensity. In addition to the expected elevation in SPL (a hallmark reflexive response to background noise), we observed systematic changes across multiple acoustic measures, including increased fundamental frequency, vowel duration, and vowel formant space expansion during noise blocks (Figure 1-S1, top row). These adjustments are consistent with previous research demonstrating that speech in noise is characterized by prosodic and articulatory modifications aimed at enhancing spectral contrast and signal robustness under masking conditions (Garber et al., 1976; Garnier et al., 2010).

Beyond the articulatory and phonatory domains, our analyses identified increased respiratory excursion during Lombard speech. Time-locked measurements revealed consistently larger respiratory excursions in noise blocks, characterized by greater dynamic range and mean amplitude in normalized respiratory traces (Figure 1–S2). These physiological changes suggest that speakers increase both inspiratory volume and expiratory drive to support elevated vocal intensity, consistent with a feedforward adjustment in respiratory planning (Stathopoulos et al., 2014). Although increased respiratory effort is a consistent feature of the Lombard effect, the specific strategies used to achieve it may vary. Winkworth and Davis (1997) observed that both increases and decreases in lung volume initiation and termination across individuals, indicating flexible rather than uniform respiratory adaptation. Our findings align with this compensatory framework, as increased excursion and dynamic range were present overall, yet the precise implementation likely differs across speakers.

Importantly, our findings are further supported by evidence that most individuals with Parkinson’s disease can effectively increase vocal intensity in noisy environments by recruiting both laryngeal and respiratory subsystems. This multisystem adaptation appears robust and can be reliably elicited by external auditory cues, such as background noise, which may trigger more natural and automatic increases in vocal intensity than explicit loudness instructions. Critically, these adaptive responses are not limited by the severity of clinical motor symptoms, as demonstrated by the lack of association between the magnitude of Lombard-induced changes and preoperative UPDRS scores or dopaminergic response (Figure 1–S4). Neither baseline motor impairment (preoperative off-medicine) nor dopaminergic response (off-medicine minus on-medicine) significantly correlated with the Lombard effect, suggesting that the underlying sensorimotor circuits remain largely preserved and can be harnessed for functional speech improvement.

Collectively, our results demonstrate that the Lombard effect engages a complex, integrated sensorimotor adaptation involving articulatory, phonatory, prosodic, and respiratory subsystems. The consistent presence of these adaptations, even during awake neurosurgery, underscores their robustness. More broadly, these findings highlight the utility of speech-in-noise paradigms for probing preserved versus compensatory mechanisms in motor speech disorders, offering promising avenues for clinical assessment and intervention.

### Ambient noise coding in basal ganglia

The basal ganglia are classically known for their role in motor control. The field has grown to appreciate that BG encoding of (multi-)sensory information—which may, in turn, subserve motor control (Chudler et al., 1995; Kameda et al., 2023; Nagy et al., 2006; Rothblat & Schneider, 1995; Strecker et al., 1985). For speech, our research group has come to view the BG as housing a ‘sensorimotor context’ for speech production at the adult stage. Our results fit into this conceptual framework, offering evidence that GP tracks ambient noise levels that can be used to modulate speech and voice parameters according to environmental constraints. This would be one of many other computational ingredients for speech production; evidence for other sensory state variables include somatosensory feedback regarding the state of the vocal tract (Chrabaszcz et al., 2019), and instantaneous auditory feedback about speech sounds (Zhu et al., 2025).

We found that ambient noise-modulated and speech-modulated populations were separate in LFP-BGA and even anti-correlated in SU-FR. What is the advantage of encoding these two variables in distinct neural populations? We speculate that ambient noise tracking could help scale movements in other motor domains of communication. Trujillo et al. (2021) found that interlocutors use changes in both the acoustic (articulatory and vocal) and visual (hand gestures and mouth movements) domains to ensure robust communication, suggesting that humans have a modality-independent energetic scaling factor that determines movement vigor (Reppert et al., 2018; Shattuck-Hufnagel & Ren, 2018). Another possible function of separate neural encoding of ambient noise would be to modulate arousal, as louder environments often induce higher states of arousal (Alvar & Francis, 2024).

Prior studies have investigated the relationship between auditory input and movement scaling (Anzak et al., 2016), but the transient nature of the auditory stimuli in those studies makes the neural and behavioral responses, in our interpretation, could be explained in terms of a ‘startle’ response (Koch, 1999, p. 19). For startle responses, transient (∼ms) responses activate the brain across many regions in unison but don’t necessarily code any stimulus-specific characteristics. The Lombard effect, and the ambient noise coding we observed in the current study, operates on a longer timescale (∼tens of seconds to minutes)—a timescale that is challenging to study in cognitive neuroscience paradigms but one that is behaviorally important and clinically relevant.

### Weak evidence for voice intensity coding in BG

Clinical and non-human primate investigation into the neural basis of motor gain or ‘vigor’ control has consistently implicated the BG (Brotchie et al., 1991; Mazzoni et al., 2007; Mink & Thach, 1991b; Turner & Anderson, 1997). Primate and human arm-reach studies have found that gamma activity in the BG correlates with movement gain (Brücke et al., 2012; Fischer et al., 2017; Joundi et al., 2012; Lofredi et al., 2018). Dastolfo-Hromack et al. (Dastolfo-Hromack et al., 2021) reported STN theta activity correlated with the second format /i/-to-/u/ ratio. We found only weak evidence of voice loudness coding in our recordings. At the cohort level, our analyses reveal no robust differences between soft, medium, and loud trials (Fig 3A, B). In LFP-BGA, there are differences between trial-averaged traces when grouped by loudness but they are only present in the mid-sentence time window, and only in the GP and not in the STN (Fig 3—S2; Wilcoxon signed-rank; p=0.0005 with all data, p=0.007 with outliers removed). The time window around speech onset shows no clear separation in either region or modality.

We propose several possibilities to explain the absence of such neural coding in our results. First, we used voice intensity as the sole measure of speech gain. Our BG speech loudness hypothesis relied on the assumption that the neural underpinnings of limb control would generalize to speech motor control. However, voice intensity control differs from limb gain control in several ways. Limb gain control is usually measured by velocity or force; voice intensity modulation, however, does not necessitate changing the speed of any articulators. The arm reach literature has found that LFP power most closely tracks movement speed. It is unclear what the analog of these parameters is for speech production.

Tan et al. (2013) found that gamma power (55-90 Hz) only correlated with effort in the most intense conditions in a manual grip task. In less intense conditions, beta was a stronger correlate of motor effort. It is possible that our experimental design did not probe the most vigorous end of speech motor gain. If gain encoding shifts from beta to gamma with increasing vigor, we should have seen an encoding in beta for speech gain.

We observed a weak pattern of loudness coding in LFP-BGA within GP, and not STN, around speech onset that could be described as an inverted U (Fig 3A). Soft speech and loud speech have lower LFP-BGA than medium-loudness speech. Other results in limb control have shown that the relationship between gamma power and movement amplitude may be an inverted U, such that the difference between medium-amplitude gamma power and low-amplitude gamma power is greater than the large-amplitude and low-amplitude comparison. We also note that Eliades et al. (2012) observed a similar pattern in single-neurons auditory cortex in response to Lombard-like masking noise in marmosets.

### Single unit-LFP relationship

Given the complexity of the relationship between LFP-BGA and SU-FR, the similarity of ambient noise coding provided strong converging evidence for the representation of ambient noise in basal ganglia activity (Fig 3). We found similar encoding of speech, noise, and speech-in-noise LFPs and single unit firing rates, albeit with different magnitudes and frequency.

In cortex, cognitive neuroscience and clinical intracranial mapping has accepted 60-150 Hz activity as a proxy for ‘multi-unit activity’ in the region of the recording electrode (Flinker et al., 2015; Lachaux et al., 2012; Mesgarani et al., 2014; Ray & Maunsell, 2011). In basal ganglia, there is less evidence and less consensus about how to interpret LFP power in the > 40 Hz range. While the spike-LFP relationship in the beta band has been studied (Bahuguna et al., 2020; Scherer et al., 2022), the >40 Hz relationship has been sparsely reported in studies. The tonic firing rate of neurons in some basal ganglia nuclei can be as high as 80 Hz (Georgopoulos et al., 1983; Hasegawa et al., 2022; Turner & Anderson, 2005), compared to that of cortical neurons under generally 10 Hz (Griffith & Horn, 1966; Roxin et al., 2011). Basal ganglia LFPs have a markedly different power spectral profile compared to cortical LFPs (Bush et al., 2023).

Inspecting time-frequency modulations at the single-channel level in our data, it was evident that increases in power ranged from ∼40 Hz to 250 Hz, and varied in bandwidth. At the grand average level, the power increases spanned 60 to 250 Hz (Fig 2--S1). Further studies are needed to define the bandwidth of modulation in GP and STN; many studies have reported power spectra using a linear scale and have failed to inspect >90 Hz activity.

Cortical LFP-BGA task-related modulations are almost exclusively in the positive direction with respect to baseline (Crone et al., 2006; Forseth et al., 2018; Lachaux et al., 2012). Increases in firing rates from low tonic FR in cortical areas is in alignment with this idea. If we take LFP-BGA as a proxy of multi-unit activity, the high tonic FR of basal ganglia nuclei, especially GP, raise the question of whether both strong positive and negative LFP-BGA deflections can be observed in these areas since both positive and negative changes in SU-FR activity are common. Our results inform this question; we observed exclusively positive speech-related deflections in LFP-BGA. This observation could at least partially be explained by the fact that a *majority* of SU-FR changes were positive during movement, leading to a net increase in LFP-BGA. The majority-increase in SU-FR in STN during speech has been a consistent finding across datasets (Lipski, 2023; Lipski et al., 2018; Zhu et al., 2025) and research groups (K. Johari et al., 2023; Tankus & Fried, 2019).

Future research in both humans and non-human primates should prioritize simultaneous recordings of SU-FR and macro-LFPs to clarify the LFP-unit relationship. Observing the relationship over the course of Parkinsonian induction in non-human primates will be particularly valuable as they will bridge the gap between basic science of single-unit neural coding and the clinically-relevant LFP scale in basal ganglia.

### Limitations

We recorded neural activity from a clinical population whose disease state confounds the generalizability of our results. Patients with Parkinson’s disease and dystonia have pathological BG activity, which could influence our results. However, PD and dystonia patients’ BG encode movement vigor in non-speech tasks (Brücke et al., 2012; Fischer et al., 2017; Joundi et al., 2012; Lofredi et al., 2018); we therefore suggest it is not solely due to disease state that we did not observe clear speech-vigor coding.

In some patients, our STN-LFP coverage was located above the dorsolateral border of the STN. A consistent trend in the LFP-BGA data was that both ambient noise and speech loudness were encoded in GP and not in STN (Fig 3, 4). We cannot completely discount the possibility that the absence of effect in STN is related to poor coverage of dorsolateral STN.

In most psychoacoustics experiments, the Lombard effect is elicited using headphones to prevent contamination of the microphone (Garnier et al., 2010). Clinical constraints of the operating room prevented us from using headphones; instead, we used a free-field approach. This, as expected, resulted in acoustic contamination on the microphone. We used acoustic echo cancellation to remove this noise, and verified with control experiments that residual leakage after the noise removal algorithm was run didn’t affect our results (Fig 1-S6).

The size of behavioral manipulations in this study were 1) subtle compared to other studies of gross motor behavior and 2) temporally imprecise in the sense that ambient noise levels changed over the course of seconds. Both of these factors make identifying neural correlates challenging.

## Methods

### Participants

Patients were approached in the weeks leading up to their awake deep brain stimulation surgery for Parkinson’s disease or dystonia at Massachusetts General Hospital. This study focused on PD and dystonia patients with STN or GPi targets, of which there were 23. Table S1 shows demographics and clinical characteristics of the population. All participants were fluent and literate in English.

All procedures were approved by the Massachusetts General Hospital Institutional Review Board (IRB Protocol #2019P003841) and all participants provided informed consent to participate in the study.

### Surgery

Target nuclei for each patient was determined by a team of neurologists and neurosurgeons at Massachusetts General Hospital based on clinical needs. Patients were taken off dopaminergic medication the night before surgery. The dorsolateral motor STN was the surgical target in STN-DBS patients. The posteroventral sensorimotor GPi was the target for GPi-DBS patients.

Surgery workflow was as follows:

1. Sedation and anesthesia
2. Headframe mounting
3. ROSA (Robotic Stereotactic Assistance) approach
4. Bilateral burr hole
5. Research electrocorticography strip insertion
6. Unilateral microelectrode mapping (3x tracks, usually medial-central-posterior in the “+” Benn-Gunn configuration) to determine track and depth at which to place DBS lead
7. Research experiments (the ‘Lombard task’ reported in this study)
8. MER removal, DBS leads inserted along determined track, at depth determined in step 6
9. DBS lead placement verified with stimulation
10. Repeat Steps 6, 8, 9 in the other hemisphere

Pre-operative MRI, intra-operative CT, and post-operative CT scans permitted localization of the microelectrode tracks used in the present study (see Electrode Localization section).

### Experimental Paradigm

Participants were supine as they engaged in a speech production experimental task in the operating room (Fig 1A). A directional microphone (Zoom SSH-6 shotgun capsule, Zoom Corporation, Tokyo, Japan) was placed ∼10 cm in front of the participant’s mouth to record produced speech. To deliver visual stimuli, a monitor was mounted to the surgical bed and arranged to face the participant at a comfortable distance (∼ 2 ft). Auditory stimuli were delivered over loudspeakers placed at the foot of the surgical bed (JBL 2P Studio Monitor).

Two experimenters administered the task: one ‘backstage’ behind the surgical drape controlling the experimental equipment and the other ‘bedside’ standing next to the patient. At the beginning of the recording session, the bedside experimenter briefly reminded the patient of what to expect during the task. Participants were exposed to the task in a pre-operative session. The bedside experimenter stood beside the participant for the duration of the session. The backstage experimenter listened to the participant’s speech over headphones and initiated the transition between trials.

Trials consisted of a 1 s intertrial gap, stimulus presentation (∼ 2 s), go cue, and speech production (∼ 2 s). The cue to start speaking (‘go cue’) was indicated by a color change in the visual stimuli; participants were instructed to wait until the text turned green before starting the utterance. After the participant completed a sentence, the backstage experimenter pressed a key on the experiment computer to initiate the next trial. We chose to make the task experimentally-timed to accommodate for the variable timing of production and to have the ability to pause or stop the task as required.

Visual stimuli were presented in white text in standard American English orthography on a black background. Sentence stimuli consisted of sentences from the *Harvard Sentences* databank (“IEEE Recommended Practice for Speech Quality Measurements,” 1969). We used prerecorded audio of each sentence from the UW/NC Corpus (Panfili et al., 2017) (speaker ID PNF133). Example sentences include: ‘Code is used when secrets are sent’ and ‘The plant grew large and green in the window’. Sentences for this study were selected from the larger set of *Harvard* sentences to include all three canonical vowels that define the vowel space in English (/i/, /a/, /u/). Table S2 lists the full set of sentences used in the task.

Trials were presented in blocks. Each block consisted of 10 trials. NOISE blocks had continuous multitalker babble playing (https://auditec.com/2015/07/02/multitalker-speech-babble-16-talkers), while QUIET blocks had no babble. Multitalker babble was played at 70 dB—calibrated for each participant using a sound-level meter (Reed R8050)—over loudspeakers. Transitions into and out of NOISE blocks consisted of a fade in/out of the babble over a 1s window. Four NOISE and four QUIET blocks (total of 80 trials) were presented in pseudorandom order; we required that two blocks of NOISE and two blocks of QUIET were presented in the first half of the experiment such that data would remain usable if a participant asked to stop the task after 40 trials.

Experimental code was written in MATLAB using Psychtoolbox (https://github.com/Psychtoolbox-3/Psychtoolbox-3). All stimuli and code for the experimental design can be found at the following Github repository: https://github.com/Brain-Modulation-Lab/Task_Lombard.

### Signal acquisition and alignment

Subcortical data were collected using the Neuro-Omega system (Alpha Omega Engineering, Nof HaGalil, Isreal) using parylene insulated tungsten microelectrodes (25 microns in diameter, 100 microns in length) with a stainless steel macroelectrode ring (0.55 mm in diameter, 1.4 mm in length) 3 mm above the tip of the microelectrode. Multiple subcortical recording locations were sampled in most participants (Table S1). LFPs were recorded on the macroelectrode, while single units were recorded from the microelectrode.

We used the Ripple neural acquisition system to record audio and respiratory signals, interface with MATLAB, send TTL (transistor-transistor logic) triggers across the system, and monitor signals in real time in the operating room (Ripple Neuro, Millcreek, Utah, USA).

To temporally align audio recording, the subcortical neural recordings, and the cortical recordings, we used TTL triggers sent from the Ripple system to the Neuro-Omega clinical system and the Zoom-H6 audio acquisition. The alignment was verified and regularly achieved sub-millisecond precision. The full code for alignment is published and available for download at https://github.com/Brain-Modulation-Lab/bml.

### Electrode localization

Participants varied in whether they were undergoing unilateral or bilateral DBS surgery, but all LFP and single-unit recordings in this study are from the left hemisphere. The locations of the micro- and macroelectrode contacts were determined using the semiautomatic approach leveraging the Lead-DBS toolbox (Horn et al., 2019). Lead-DBS localizes the implanted DBS probe by coregistering the native-space pre-operative MRI with the post-operative CT scan, which contains the electromagnetic field distortion artifact of the DBS probe’s contacts. The post-operative CT was co-registered to the pre-operative MRI using a two-stage linear registration (rigid followed by affine) as implemented in Advanced Normalization Tools (Avants 2008; http://stnava.github.io/ANTs/). Because the DBS probe followed one of the three (medial/lateral, central, posterior) tracks tested during the surgery and was implanted at a known depth relative to the intraoperative MER recording probes, we could infer the locations of the MER probes within the MRI volume. After localization and defining atlases in native space, Lead-DBS computes a mapping between native space and Montreal Neurological Institute MNI152 NLIN 2009b space (explained further here: <u>WebArchive</u> https://www.lead-dbs.org/about-the-mni-spaces/). These MNI coordinates were used in visualizations of recordings sites (Fig 2). STN and GPi/GPe surfaces were used from the DISTAL atlas (Ewert et al., 2018).

See the script preproc/sub_DM_example_ses_intraop_B04_mer_localization.mlx in the project repository for full details of the localization process.

#### Acoustic Echo Cancellation

Experimental paradigms studying the Lombard effect often use headphones to deliver multitalker babble to participants as they speak (Lu & Cooke, 2008; Stathopoulos et al., 2014), despite the aural occlusion confounding the generalizability of these studies to real-world scenarios (Vaziri et al., 2019). Headphones were not used in this study due to the complex setup of neurosurgery in the operating room and ethical reasons to maximize communication between patients and the medical team. Instead, we delivered the stimuli and background noise multitalker babble over loudspeakers. There was, as a result, background noise contamination in the directional microphone that was intended to capture the participant’s voice. We used Acoustic Echo Cancellation (AEC) to remove this contamination, ensuring that changes in voice intensity were indeed from the participants’ voice intensity changes and not from background noise intensity changes. Directional microphone .wav files along with stimulus .wav files recorded synchronously were processed with the SpeexDSP Python (https://gitlab.xiph.org/xiph/speexdsp). See the full code for AEC in the script wav_aec.py in the paper code repository.

We validated the AEC procedure on the first 12 patients who completed the task. Audio was calibrated with 3 beeps of known intensity (recorded on the sound-level meter) at the beginning of each task run. The mean intensity of the beeps was 73.1 dB. The mean SPL was 67.6 ± 4.3 dB in the quiet condition and 70.9 ± 4.0 dB in the noisy condition. The mean increase in vocal intensity from the quiet to noisy condition was 3.6 dB. The mean SPL for the intertrial gap in the quiet condition was 51.7 dB prior to applying the AEC algorithm and 49.2 dB afterward, with a 2.6 dB difference. In the noisy condition, the mean SPL for the intertrial gap was 64.5 dB prior to applying the algorithm and 52.4 afterward.

#### Phonetic Transcription

To annotate participants’ speech in each trial, we used a semi-automated approach: automated forced alignment followed by manual verification. Produced speech audio waveforms were extracted for each trial. We then force-aligned the audio with the Penn Phonetics Forced Aligner (Yuan & Liberman, 2008) using the English orthography for each sentence. A trained Speech-Language Pathologist manually verified—and edited if necessary—the alignment using Praat (Boersma & Weenink, 2007). If the forced alignment failed, we iteratively took smaller and smaller chunks from the waveform until the alignment succeeded. Speech-Language Pathologists qualitatively indicated speech dysfluencies and errors in each trial. Speech onset in each trial was defined as the onset time of the first phone, and speech offset was the the offset time of the last phone.

#### Voice Intensity Estimation

Voice intensity was measured in Praat. Audio waveforms for each run were passed to Praat. Normalized intensity curves were extracted using a command line call to a Praat script, which implemented a call to ‘Sound: To Pitch…’ which extracts both pitch and intensity estimates. We specified a pitch floor and pitch ceiling of 100 and 500 Hz for female participants. For male participants, the values were set to 60 and 300 Hz. See https://www.fon.hum.uva.nl/praat/ for further details regarding estimation of intensity.

Analyses were repeated using the waveform RMS at each time point, yielding similar results.

#### Neural Signal Processing

##### Local Field Potentials

LFPs were stored in FieldTrip ‘raw’-type objects after low pass filtering at 250Hz using a 4th order Butterworth filter, and downsampling to 1KHz and stored as a Fieldtrip object. We applied a 5th order high-pass Butterworth filter at 1 Hz to remove drifts and low-frequency components.

Broadband gamma activity (BGA) was extracted trial-by-trial after excluding trials that were deemed artifactual. We used multitaper Discrete Prolate Spheroidal Sequences (DPSS) time-frequency analysis to maximize signal-to-noise ratio in BGA. The algorithm was implemented in FieldTrip ft_freqanalysis() with the following parameters at 10 ms strides: {method=”mtmconvol”, taper=”dpss”, foi=130, tapsmofreq=65}. The power of each complex timeseries was returned. For each run for a given subject, this resulted in a tensor of shape n_trials x n_channels x n_timepoints. Data tensors were later reshaped and concatenated across all runs and subjects for cohort analyses in R. See the script 20230524-subctx-lfp-group-PLB/A01_prepare_subject_TFR.m and the function bml_extractgamma() for full recapitulation of our BGA extraction pipeline.

BGA values in the paper can be interpreted as power in the 60-200 Hz frequency band. BGA values were log-transformed to make them more Gaussian and therefore more amenable to statistical analysis. BGA time-series were not immediately normalized with respect to baseline (e.g., via z-score) because we noted non-stationarities over the course of runs in the signals (Fig 2--S2). Instead, we decided to explicitly model the non-stationarities in our statistical approach (see Statistics section below).

The spectrograms in Fig 2-S1 were calculated with 7-cycle Morlet wavelets with 40 log-spaced filter banks between 4 and 250 Hz. Power estimates at each time were baselined to the [-1, 0] s window before stimulus onset.

##### Single units

Subcortical microelectrode signals were filtered with a zero-phase lag filter in the 300- to 3,000-Hz band, and spikes were detected and sorted using the semiautomated template-matching algorithm OSort (v4.1) (Rutishauser et al., 2006b). To identify putative single units and assess their quality, we computed several spike sorting quality metrics (mean ± sd, Figure 2—S1): (i) the percentage of interspike intervals (ISIs) below 3 ms was 1.71% ± 1.88%; (ii) the ratio between the standard deviation of the noise and the peak amplitude of the mean waveform of each cluster was 12.69 ± 2.53 dB (peak SNR); (iii) the pairwise projection distance in clustering space between all neurons isolated on the same wire was 11.24 ± 6.39 (projection test (Pouzat et al., 2002); in units of standard deviation of the signal); (iv) the modified coefficient of variation of variability in the ISI (CV2) was 0.84 ± 0.14; and (v) the (log) isolation distance (Harris et al., 2000) was 1.26 ± 0.37. We calculated the isolation distance in a ten-dimensional feature space (energy, peak amplitude, total area under the waveform and the first five principal components of the energy-normalized waveforms). These metrics were used to grade the quality of the putative single-units: A (stationary single-units), B (stationary units with multi-unity contamination), C (non-stationary single-units) and D (non-stationary units with multi-unity contamination).

### Data selection

#### Local field potentials

Trials in each channel were rejected first in a round of automatic artifact detection, followed by manual verification and rejection. Segments with conspicuous high-power artifacts were identified using an automatic data cleaning procedure, based on a power-based threshold. The automatic rejection was a simple amplitude-based time window rejection; line-noise-filtered signals were binned into 1 s nonoverlapping segments. We calculated the maximum absolute amplitude in each segment and log-transformed the data. By default, we rejected segments exceeding 2.5 std (∼10-fold higher the mean). After visualizing the histogram of binned amplitudes, we increased the rejection threshold if the data followed a normal distribution, as this often indicated a particularly clean data collection. Trials with time segments flagged as artifactual were discarded and channels with more than 30% of artifactual time bins were not included in the analysis.

#### Single-units

Single units of all quality categories were used in initial analyses. However, units with less than 10 trials in either the NOISE or QUIET condition were excluded.

### Time Warping

We used time warping to align behavioral events across trials. We defined the following events of interest: stimulus onset (time = 0 s), stimulus offset (t = 2 seconds), speech onset (t = 3 s), and speech offset (t = 5 s). The timestamp of each event was identified and warped to the prescribed times above. The time in any given trial, then can be thought of as a percentage complete for given time window; for example, if a participant’s speech on a given trial occurred between 3.3 – 5.3 s relative to t=0 stimulus onset, then the time 4.3 s in the trial would be considered 50% of the way through speech production; and therefore mapped to 50% complete in the time-warped data, to t = 4 s, since 50% complete of the [3, 5] s window is 4 s.

### Statistics

Statistical analyses and visualizations were implemented in MATLAB and R (R Core Team, 2017) using Tidyverse packages (Wickham et al., 2019). LFP-BGA and SU-FR activity were stored in data tables downsampled to 100 Hz. To unify timepoints across channels and smooth neural traces, time series were binned into 50 ms bins and smoothed with a 3^rd^-order, 51-point Savitsky-Golay filter implemented in R (signal::sgolayfilt()).

#### Behavioral Analyses

We calculated mean intensity values in the BSLN and SPEECH window for each trial by epoching intensity signals according to behavioral annotation tables. Cohort analyses consisted of paired t-tests using 2P total data points, where P is the number of participants in the cohort, as each participant corresponded to 2 data points: the mean intensity in the NOISE condition, and the mean intensity in the QUIET condition. Within-participant unpaired t-tests were performed to validate the effect within individual subjects; each test consisted of T_noise+T_quiet data points, where T_noise is the number of NOISE and QUIET trials the participant performed on a given run.

#### Non-stationarities in neural signals

Visualizations of absolute LFP-BGA and SU-FR revealed distinct non-stationarities in the subcortical recordings (Figure 2--S2). Over the course of minutes, the LFP-BGA and SU-FR could change by more than 40% in absolute magnitude. Non-stationarities are commonly observed immediately after electrode implantation. Due to the time constraints of intraoperative research, it was not possible to wait for the signals to stabilize. We could not, therefore, make the common assumption in cognitive neuroscience that the baseline trial window was stationary. To ensure that these non-stationarities did not drive spurious effects in the task-related modulations we were interested in, we both a) modeled slow non-stationarities using time and time-squared terms as regressors and b) ran permutation tests.

#### Determining significant GP, STN, and GP vs STN activation

Significant divergence between GP and STN time series traces was determined using across-channel bootstrapping at each time point relative to STIMULUS offset (Fig 2D, ‘Activation Timecourse’). We bootstrapped a GP-minus-STN distribution at each time point by sampling one GP channel mean and one STN channel mean and taking their difference.

We estimated the peak of activation using within-channel linear models followed by across-channel bootstrapping. To do so, we used downsampled LFP-BGA and SU-FR traces to 100 Hz. For each channel, we ran sliding window unpaired t-tests comparing average activity within a 250 ms window with respect to baseline. We tested time windows every 100 ms from -2 s to 2 s relative to speech onset. For each channel, we then extracted the time of peak activation by taking the argmax of the beta-coefficients-over-time from the t-tests. We bootstrapped distributions of time-to-peak in STN and GP separately (n=500 samples). We then bootstrapped a GP-minus-STN distribution by sampling a channel from GP and a channel from STN and taking the difference in each bootstrap draw (n=500 samples). Significance was determined by these distributions’ lower (2.5%) and upper (97.5%) bounds.

#### Permutation tests to identify linear models’ significant coefficients

We used residual permutation tests to assess the significance of regression coefficients at each recording channel. Let neural activity from a given trial and time window be denoted as A. We fit a multiple linear regression model, M₁, of the form

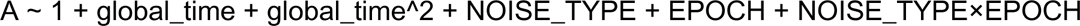

where:

- global_time + global_time^2 represent the linear and quadratic terms of solar time (capturing slow drifts or curvature over the course of a run),
- NOISE_TYPE is a categorical variable coding the background condition (NOISE or QUIET),
- EPOCH is a categorical variable coding task period (SPEECH or BSLN), and
- NOISE_TYPE×EPOCH represents their interaction term.

This model yields coefficients for the effects of time, background noise, speech epoch, and their interaction.To determine whether a given coefficient (for example, the noise main effect) was significantly different from chance, we used a residual permutation test. First, we fit a reduced model excluding the factor of interest (e.g., without the NOISE_TYPE term) to obtain predicted values A_M0 and residuals A_{res, M0}. Residuals were shuffled in a way that respected the experimental design (i.e., shuffled within noise blocks, then the block order was permuted). The shuffled residuals were added back to the model predictions to create permuted data: A_shuff = A_M0 + shuffle(A_{res, M0}). We then refit the model with all predictors:

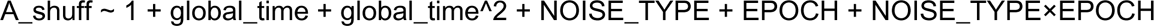

The resulting coefficient for the shuffled dataset (beta_{NOISE, null}) was stored, and this residual-permutation procedure was repeated 1,000 times to build a null distribution of coefficients.A coefficient was deemed significant if its observed value fell outside the [2.5, 97.5] percentile interval of its null distribution (e.g., Fig. 3C, insets).

By shuffling only the residuals of the null model but then running the full model, this procedure maintains the structure of the model and offers a conservative approach to estimating coefficient significance.

For visualizations, we used the residuals of the model A ∼ 1 + time + time^2, where A represents LFP-BGA or SU-FR neural activity and time is the time in seconds since the start of the run.

#### Spatial clustering

We aimed to determine if speech-active recording sites were spatially aggregated relative to all recording sites with a shuffle permutation test. We tested whether the physical pairwise distances amongst the set *speech-active* recording sites significantly was smaller than the distances amongst the set of *all* recording sites. We generated a null distribution of distances between all recording sites within a target (either STN or GP) by bootstrapping the average Euclidean distance between sites, and repeating 500 times. We then did the same after filtering the data to include only speech-active sites. Speech-active sites were considered ‘clustered’ if the actual average pairwise distance between the speech-active sites was smaller than the lower bound of the 95% CI of the null distribution.

#### Demixing PCA

We implemented the demixing-PCA pipeline described in the original paper (Kobak et al., 2016) (https://github.com/machenslab/dPCA) on our data. Time-warped LFP-BGA and SU-FR data in tabular format from Fig 2, 3 were binned into 100 evenly-spaced time bins and averaged within bin. Data were then compiled into a 4-d tensor representing [channel, condition={QUIET, NOISE}, time bin, trial number]. This is the type of tensor that dPCA accepts as input. We used standard parameters with the exception that we used a wider range of regularization lambda values, from 1e-7 to 100. For SU-FR data, dPCA found lambda=6.95e-5 maximized cross-validation performance. For LFP-BGA, the optimal lambda was 0.0483. The full code for dPCA analyses can be found at 20230524-subctx-lfp-group-PLB/A06_dpca.m

## Terminology, Definitions, and Abbreviations

GP: globus pallidus
STN: subthalamic nucleus
BG: basal ganglia
LFP-BGA: local field potential broadband gamma (70-250 Hz) activity
SU-FR: single unit firing rate

## Data and Code Availability

Data will be made available at request to the corresponding author. Preprocessing and figure-generating code is available at https://github.com/Brain-Modulation-Lab/Paper_Lombard-ambient-noise-GP-STN/

## Acknowledgements

We express our gratitude to all the patients who participated in this study.

We thank Andrew Meier from the Boston University Speech Neuroscience Lab for significant feedback on this manuscript.

## Contributions

CRediT (Contributor Roles Taxonomy) statement: Latane Bullock LB; Matteo Vissani MV; Alan Bush AB; Jackie S Kim JSK; Lori L Holt LLH; Julie Fiez JF; Robert S Turner RST; Todd M Herrington TMH; Jeffrey Schweitzer JS; Frank H Guenther FHG; Robert Mark Richardson RMR

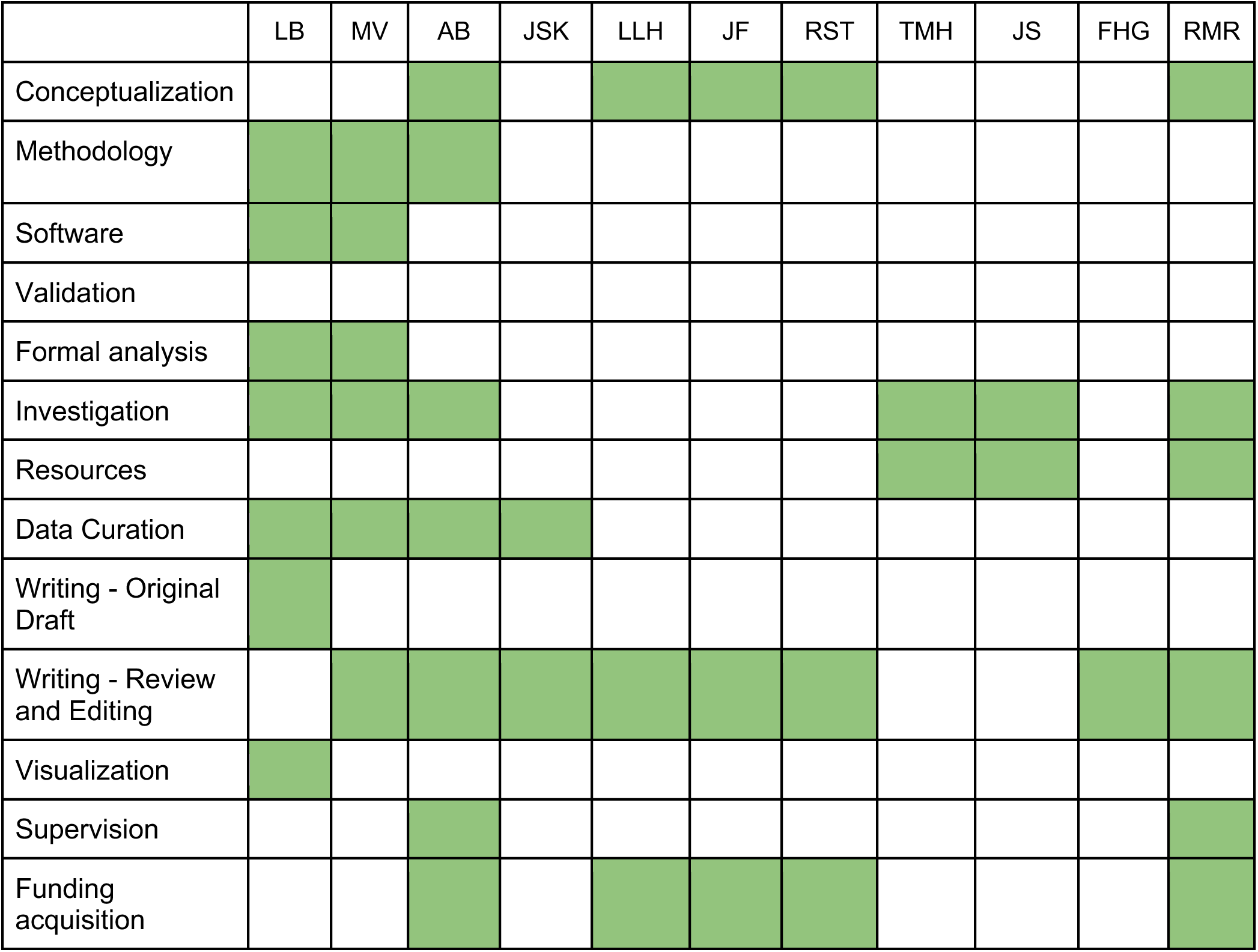

---

**Figure 1-S1:**
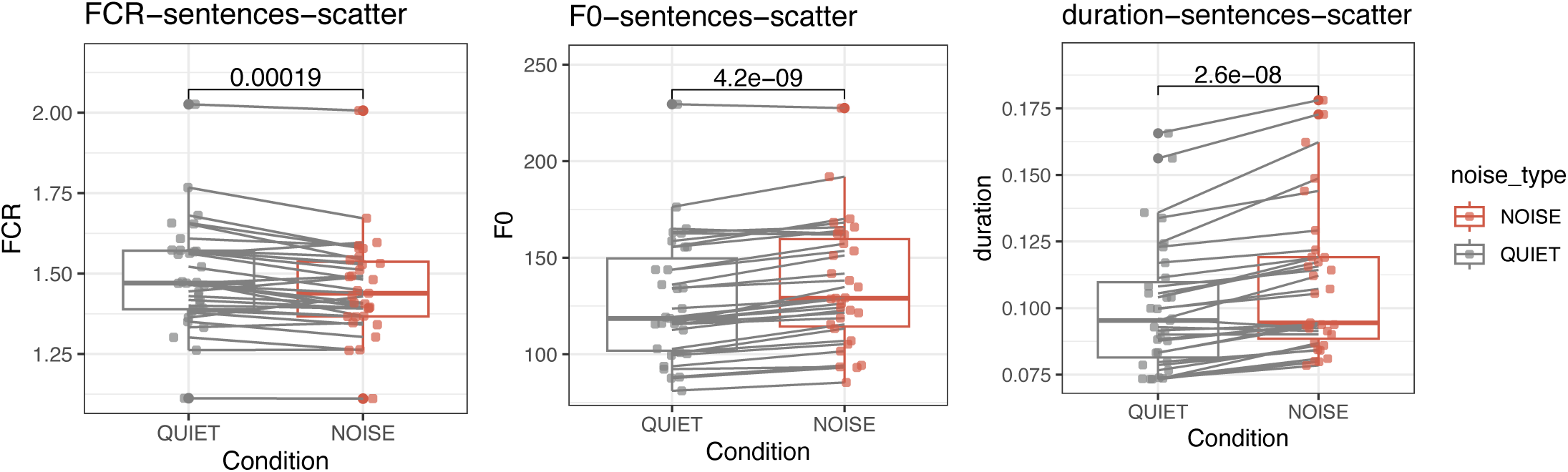
Other measures of speech gain: FCR, F0, vowel duration. Formant centralization ratio (FCR; left), fundamental frequency F0 (middle), and vowel duration (right) in the NOISE vs QUIET conditions. Each point represents the mean value across trials for each subject. FCR decreased (Wilcoxon signed-rank, p=0.00019), while F0 (p=4.2e-9) and vowel duration (p=2.6e-8) increased.

**Figure 1--S2:**
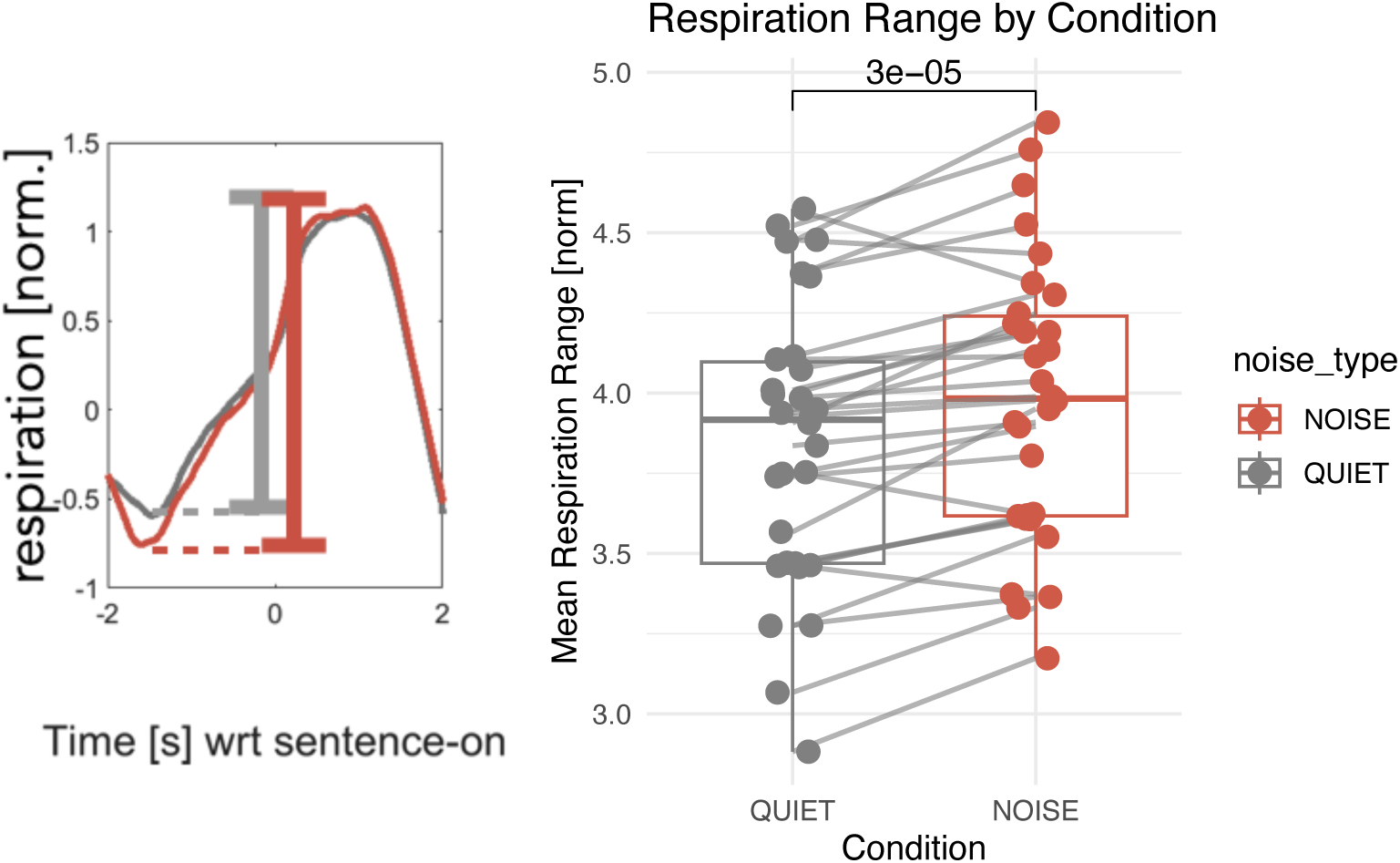
Participants breathed deeper in the NOISE condition. Respiration depth as measured by a force transduction belt around participants’ chest. Respiration traces were z-scored across the recoring session and time-locked to sentence onset. Minimum and maximum values in the [-2, 2] second window around speech onset for each sentence. The absolute different between the two defined a range for each sentence. (Left) Example traces from one subject in the NOISE (red) and QUIET (grey) condition. (Right) Cohort-wide comparison of NOISE and QUIET conditions. Each point represents the mean range for a subject in the given condition. Respiration range was higher in the NOISE vs QUIET conditions (Wilcoxon signed-rank, p=3e-5)

**Figure 1-S3:**
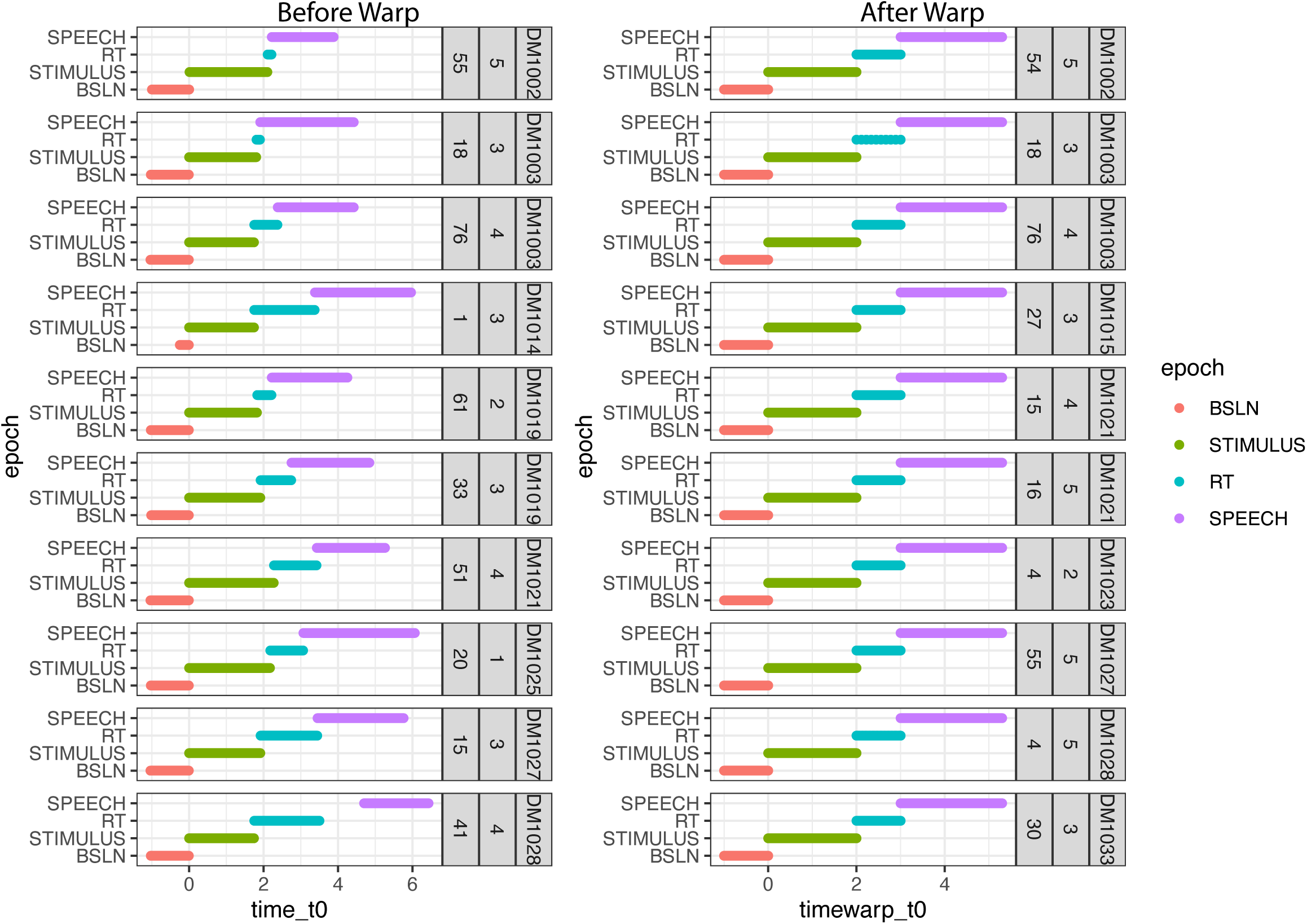
Subject and trial behavioral variability. Behavioral variability across trials. (Left) Trial time annotations for a random selection of trials across the cohort before warping. (Right) Trial time annotations after linear piecewise warping was applied.

**Figure 1-S4:**
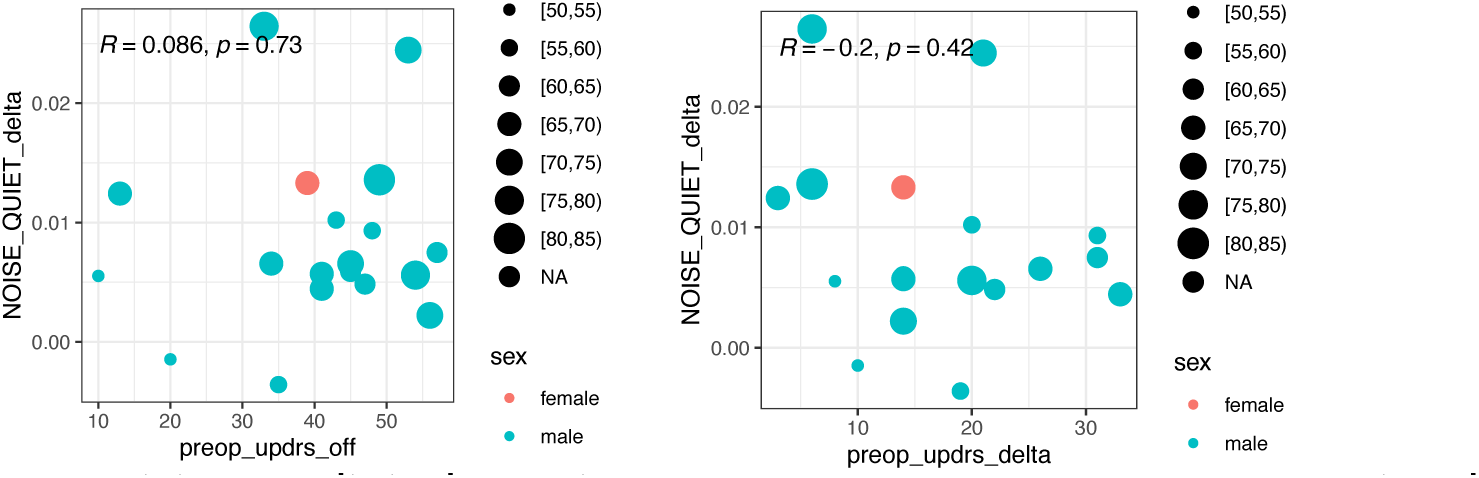
Clinical severity does not predict Lombard effect size. Relationship between participants clinical severity and the magnitude of speech-in-noise changes. We defined the ‘Lombard effect size’ to be the difference between the intensity of seech in the NOISE condition vs that in the QUIET condition. Clinical severity was approximated by the pre-operative UPDRS total score (left). No significant relationship was observed (Spearman, R=0.086, p=0.73). Plots indicate sex (color) and age (size) of participants. We also tested measured clinical severity by the magnitude of relief confered by the operation (right). No relationship was observed (Spearman, R=-0.2, p=0.42). Sex (color) and age (size) are indicated for each participant.

**Figure 1-S5:**
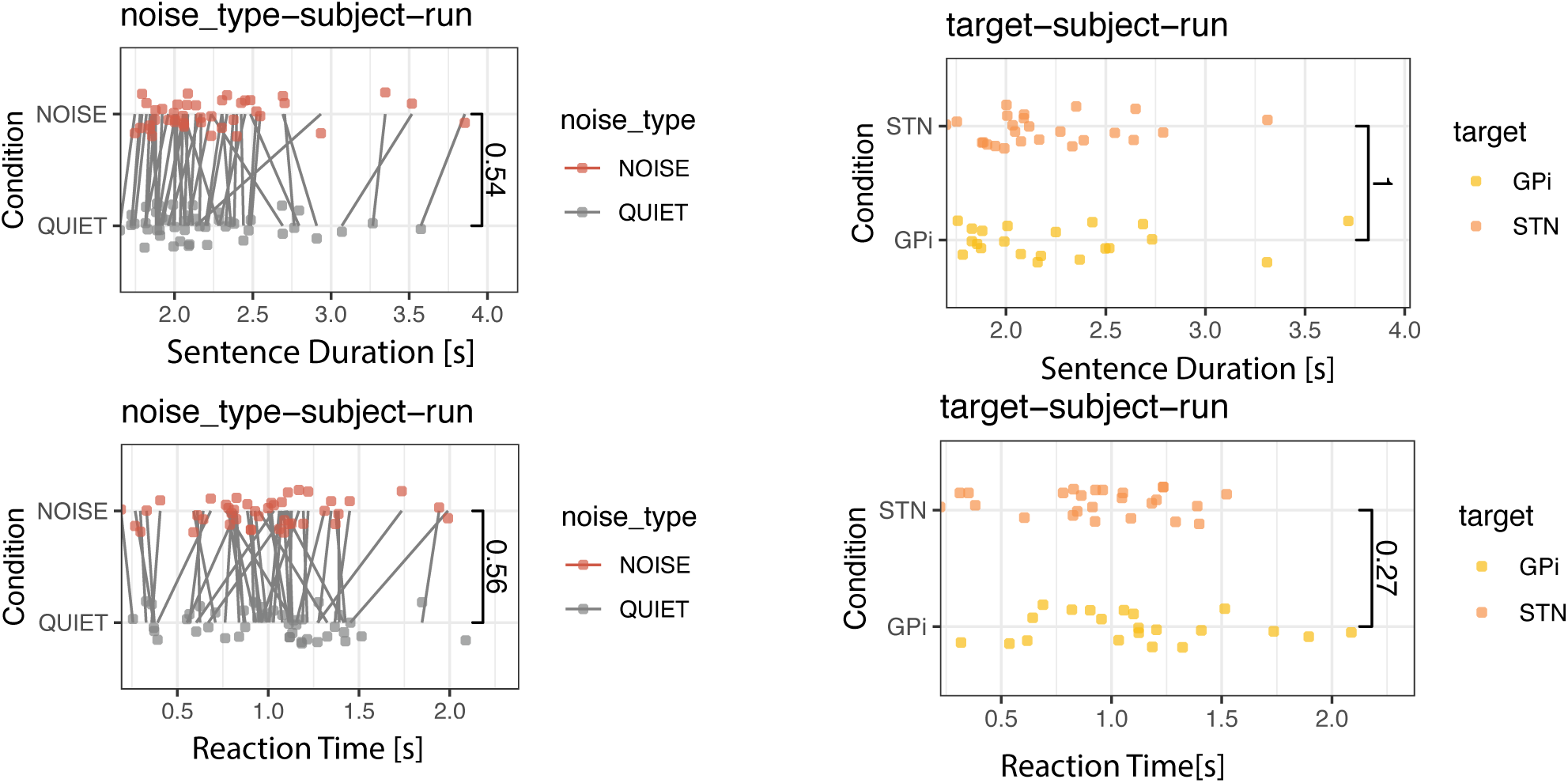
No difference in response times, sentence duration in NOISE vs QUIET, STN vs GPi target. Coarse comparison of sentence duration and reaction time between conditions (left panels) and and between deep brain stimulation targets (right panels). Each point represents the mean value across trials for each subject. No significant relationships were observed.

**Figure 1--S6:**
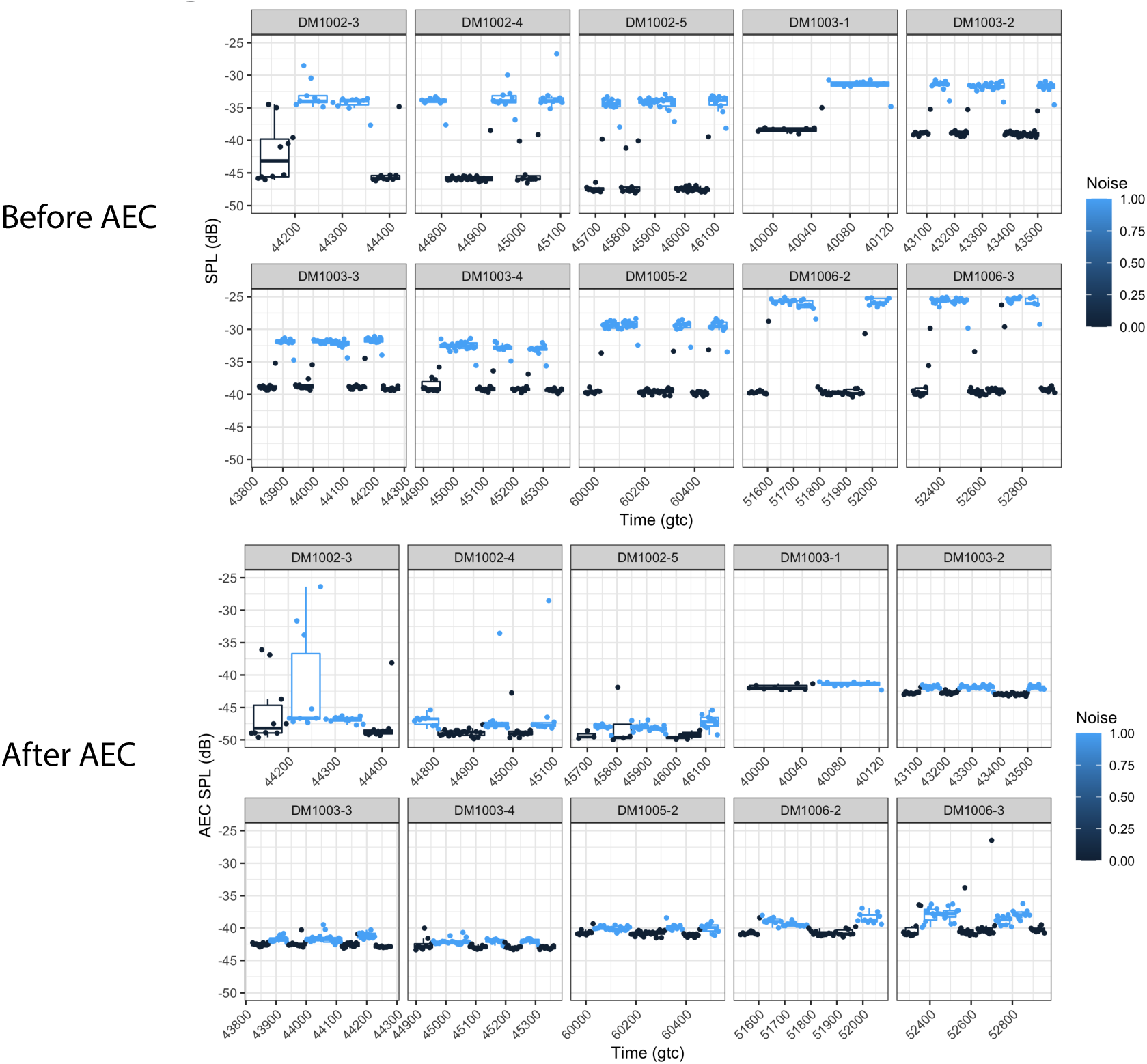
Acoustic Echo Cancellation Controls.

**Figure 2--S1:**
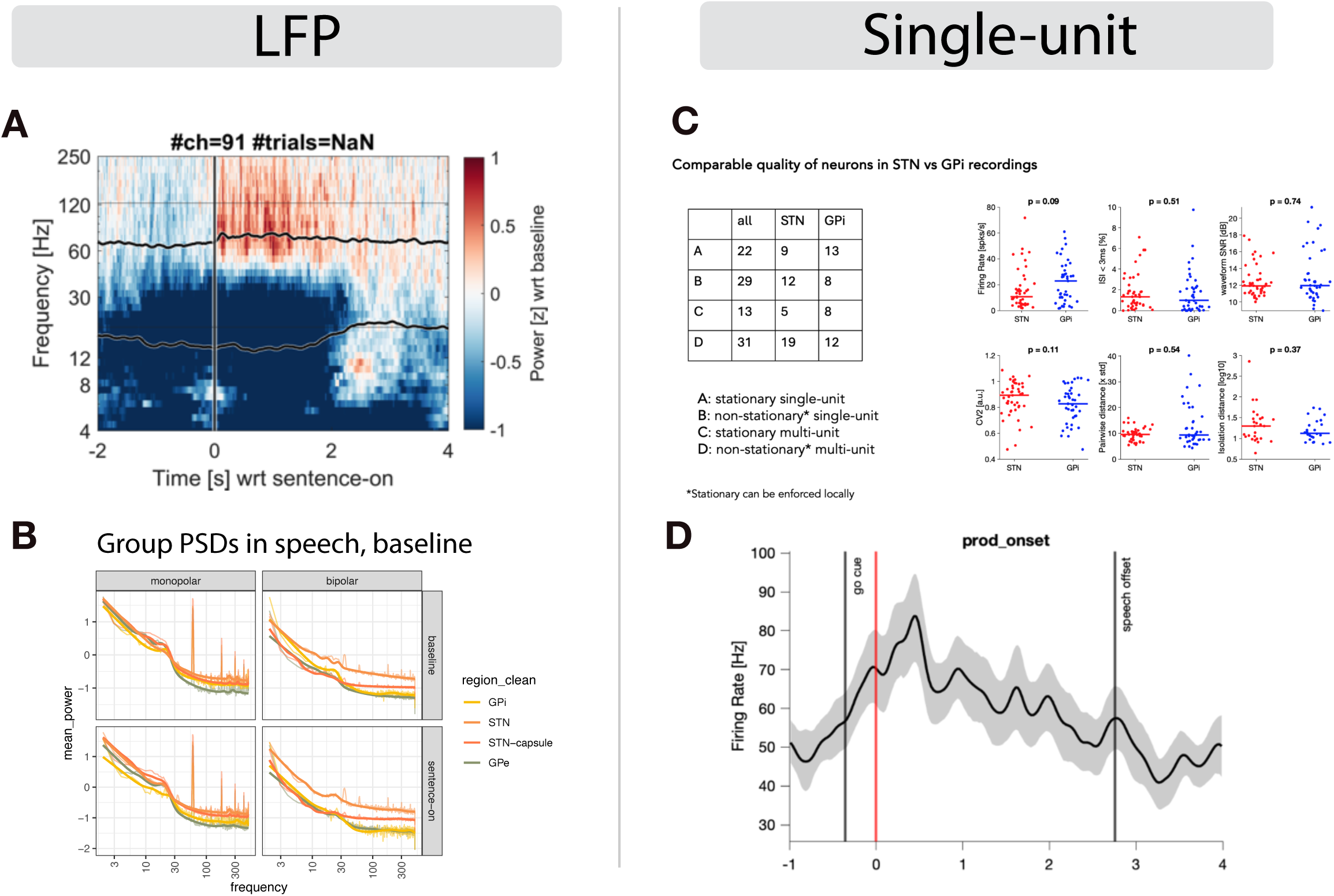
Quality metrics and example units. (A) Grand average (91 channels, across all subjects) LFP power with respect to baseline time-locked to speech onset. Trials were averaged within channel, and then channels were averaged together. Each frequency band is z-scored with respect to baseline. Black traces represents averaged power in beta (12-30 Hz) and broadband gamma (60-250 Hz). Red denotes an increase in power, blue denotes decrease. Average speech offset is 2.3 seconds. (B) Power spectral densities (thin lines) and there parameterized approximations (thin lines), averaged across all recordings and subjects. (C) Single-unit quality in STN and GPi. (D) Example speech-modulated unit. Time 0 represents speech onset. For the example subject, average speech offset was ∼2.75 s.

**Figure 2--S2:**
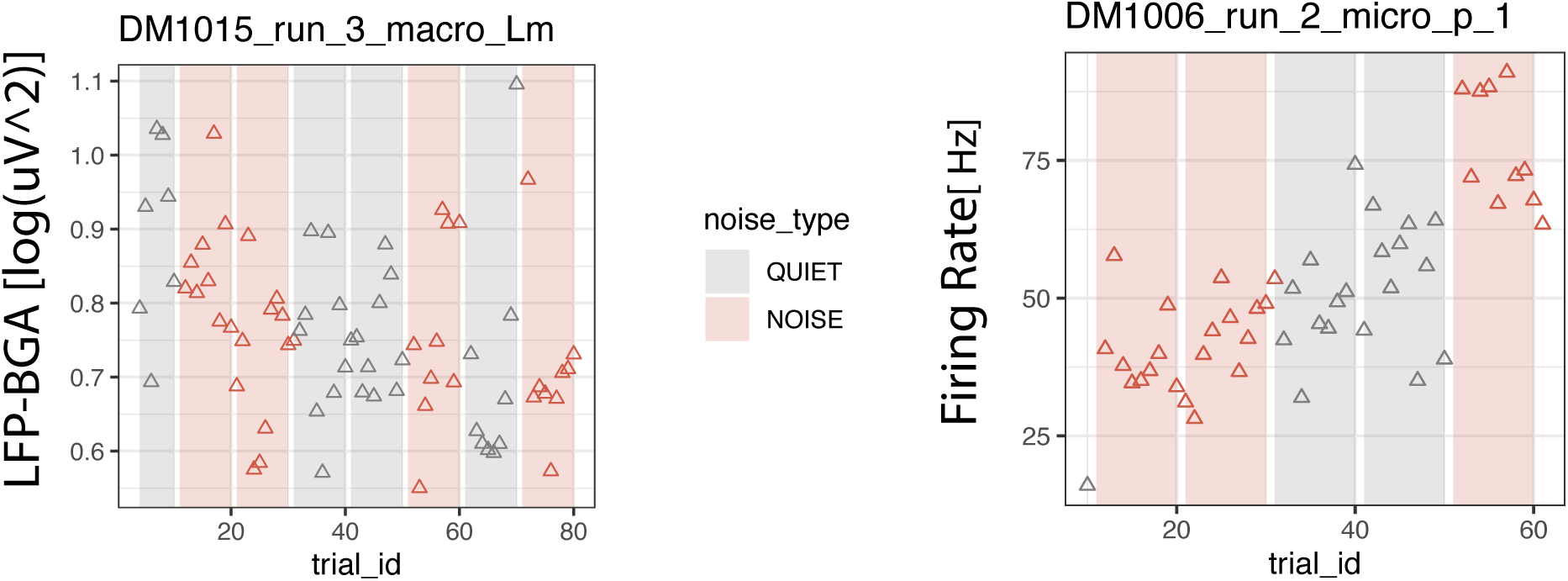
Non-stationarities in neural signals. Power (for LFPs, left) or firing rate (for SUs, right) in the baseline window of each trial in two sample subjects. Note the non-stationarities; LFP-BGA power decreased over the course of 80 trials (∼15 minutes) at the selected recording site, and SU-FR increased over the course of 60 trials in the selected neuron.

**Figure 3--S1:**
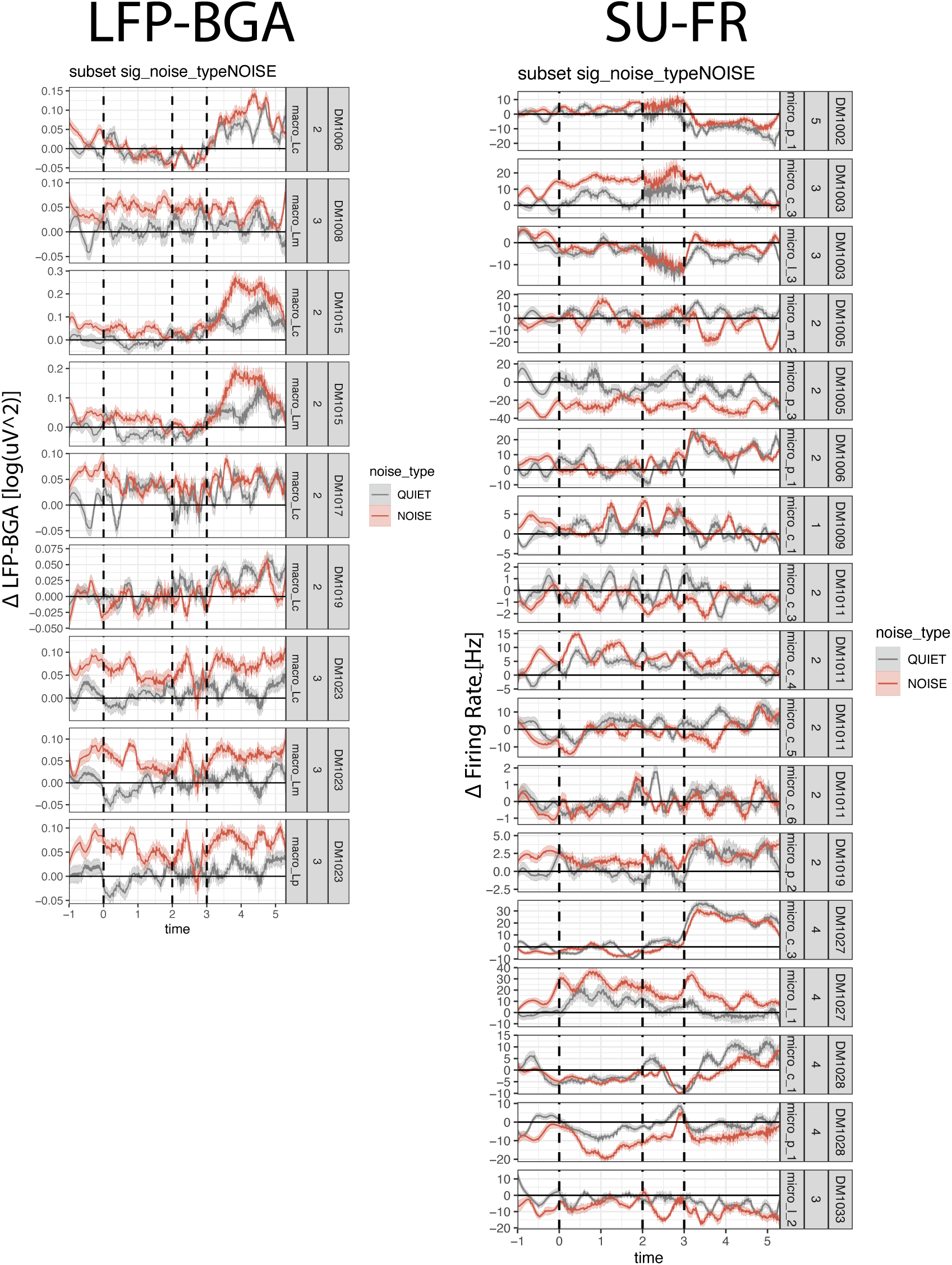
Example ambient-noise modulations. Traces of individual channels (LFP-BGA) or units (SU-FR) that were identified as being ambient noise modulated. Each plot contains a timeseries (mean ± SEM) for either LFP-BGA (left) or firing rate (right) with respect to baseline.

**Figure 3--S2:**
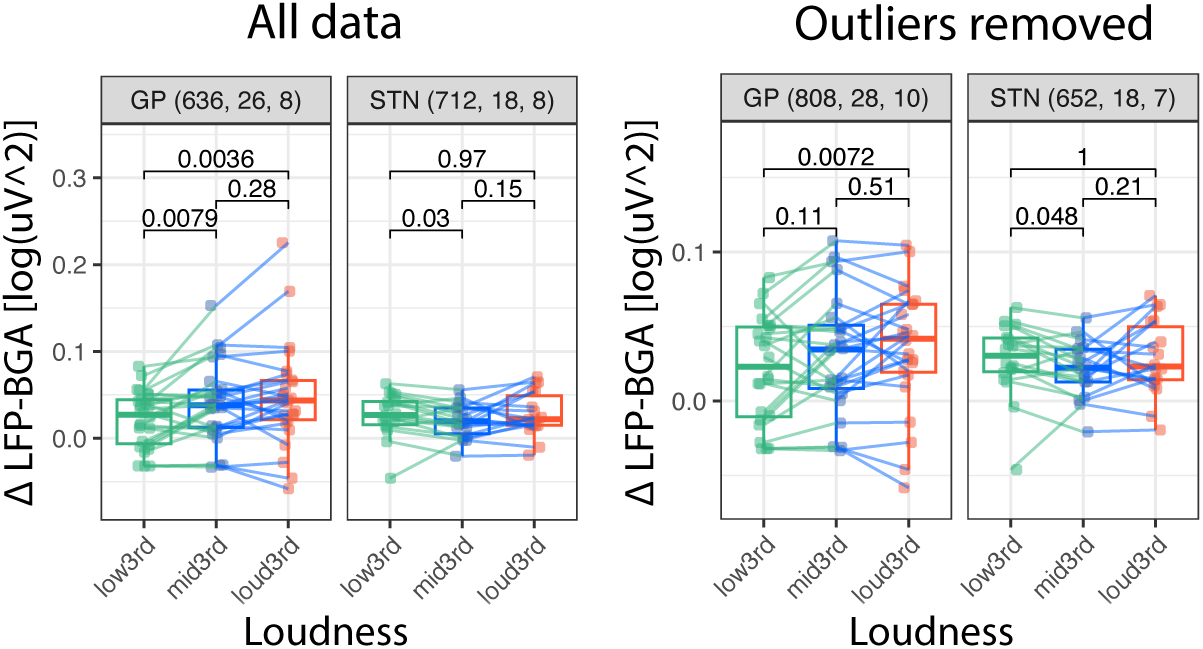
Weak coding of speech-in-noise in LFP-BGA mid-sentence. LFP-BGA with respect to baseline in the mid-speech window: [1, 2] seconds after speech onset. Each point represents a macro-LFP recording site. Data are grouped by produced speech loudness level (lowest/softest third, middle third, and loudest third) defined within each participant. Pairwise Wilcoxon signed-rank tesvvt p-values are indicated between groups. Note the significant loud-vs-soft comparison in GP (p=5.3e-4) that is preserved even when outliers (beyond mean ± 1.5 x interquartile range) are removed (p=7.2e-3).

**Figure 3--S3:**
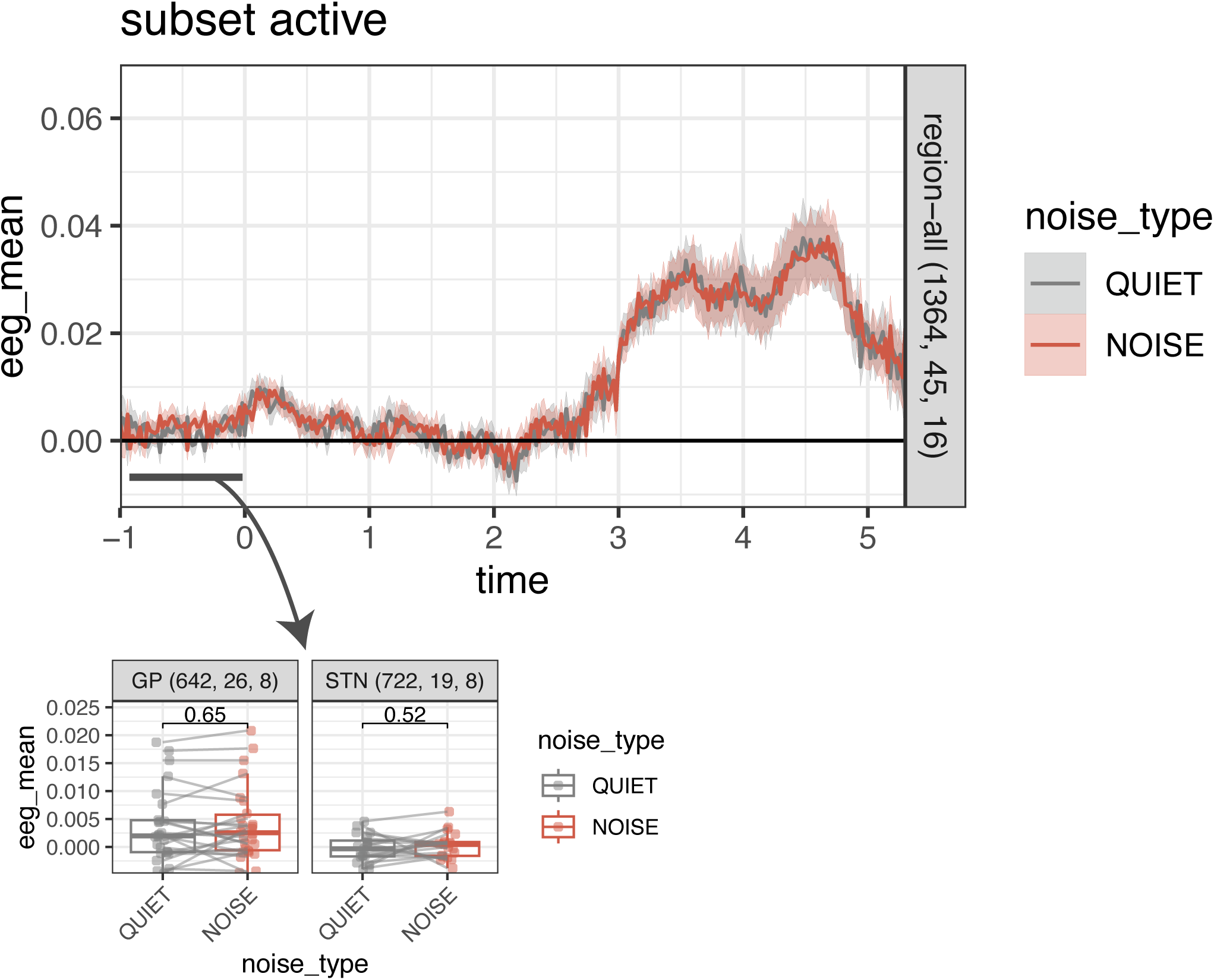
Ambient noise effect is eliminated with shuffled NOISE labels. Same as Fig 3C but computed on data with shuffled noise_type labels. The group NOISE > QUIET effect is abolished.

**Figure 4--S1:**
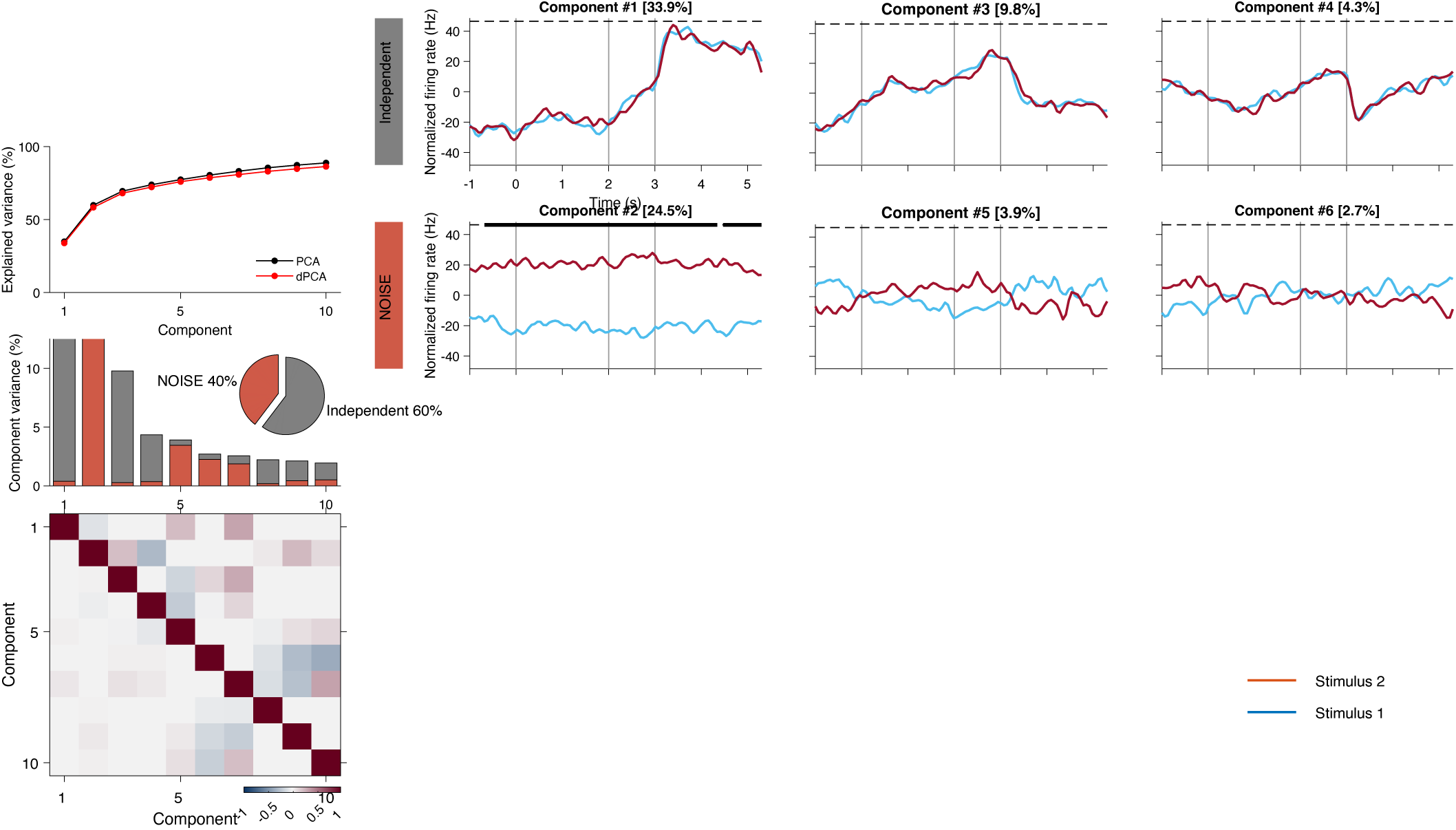
dPCA SU-FR without stationarity correction. Legends as in Figure 4 but with firing rates that have no non-stationarity correction applied. Note the exaggerated ambient noise coding in Component 2.

