## Supplemental Tables for "Independent Basal Ganglia Neural Populations Encode Speech Production and Ambient-Noise Levels"

| subject_id | age | sex | diagnosis | target | preop_updrs_off | preop_updrs_on | handedness | # SU | # SU runs | # SU trials | # LFP chans | # LFP runs | # LFP trials |
|---|---|---|---|---|---|---|---|---|---|---|---|---|---|
| **DM1002** | 82 | male | PD | STN | 49 | 43 | Left | 6 | 3 | 150 | 9 | 3 | 109 |
| **DM1003** | 59 | male | PD | STN | n/a | NA | Left | 7 | 2 | 158 | 9 | 3 | 182 |
| **DM1005** | 16 | female | Dys | GPi | n/a | n/a | Right | 7 | 1 | 79 | 3 | 1 | 75 |
| **DM1006** | 61 | female | Dys | GPi | n/a | n/a | Right | 5 | 2 | 108 | 6 | 2 | 124 |
| **DM1008** | 61 | female | Dys | GPi | n/a | n/a | Left | NA | NA | NA | 3 | 1 | 68 |
| **DM1011** | 67 | female | PD | STN | 39 | 25 | Right | 6 | 2 | 156 | 4 | 2 | 143 |
| **DM1012** | 73 | male | PD | GPi | 56 | 42 | Right | 1 | 1 | 65 | 3 | 1 | 47 |
| **DM1013** | 52 | male | PD | GPi | 20 | 10 | Right | 2 | 1 | 10 | 3 | 1 | 46 |
| **DM1014** | 58 | male | PD | STN | 35 | 16 | Right | 2 | 1 | 33 | 1 | 1 | 18 |
| **DM1015** | 64 | male | PD | GPi | 47 | 25 | Right | 2 | 1 | 31 | 4 | 2 | 144 |
| **DM1017** | 53 | male | PD | GPi | 10 | 2 | Right | NA | NA | NA | 2 | 1 | 46 |
| **DM1018** | 74 | male | PD | STN | 45 | n/a | Right | 2 | 1 | 79 | 2 | 1 | 63 |
| **DM1019** | 74 | female | PD | GPi | n/a | NA | Right | 12 | 2 | 119 | 9 | 3 | 142 |
| **DM1020** | 77 | male | PD | STN | 54 | 34 | Right | 5 | 1 | 28 | 3 | 1 | 66 |
| **DM1021** | 59 | male | PD | GPi | 43 | 23 | Right | 1 | 1 | 15 | 3 | 1 | 48 |
| **DM1022** | 55 | male | PD | STN | 48 | 17 | Right | 3 | 2 | 75 | 6 | 2 | 158 |
| **DM1023** | 62 | male | PD | GPi | 45 | 31 | Right | 2 | 2 | 55 | 6 | 2 | 112 |
| **DM1027** | 69 | male | PD & ET | STN | 41 | 8 | Right | 5 | 2 | 156 | 6 | 2 | 158 |
| **DM1028** | 59 | female | PD | GPi | NA | NA | Left | 3 | 1 | 79 | 3 | 1 | 79 |
| **DM1029** | 68 | male | PD | STN | 41 | 27 | Right | 5 | 2 | 109 | 6 | 2 | 158 |
| **DM1033** | 62 | male | PD | STN | 57 | 26 | Right | 2 | 1 | 59 | 3 | 1 | 77 |
| **DM1034** | 79 | male | PD | STN | NA | NA | Right | 3 | 1 | 74 | 3 | 1 | 57 |
| **DM1037** | 69 | male | PD | STN | 34 | 8 | Right | 2 | 1 | 55 | 3 | 1 | 79 |

Table S1. Patient demographics and data quantities. SU=single unit. Chans=channels

| sentence_id | sentence |
|---|---|
| 1 | GLUE THE SHEET TO THE DARK BLUE BACKGROUND |
| 2 | THERE ARE MORE THAN TWO FACTORS HERE |
| 3 | THE PLANT GREW LARGE AND GREEN IN THE WINDOW |
| 4 | FILL THE INK JAR WITH STICKY GLUE |
| 5 | CODE IS USED WHEN SECRETS ARE SENT |
| 6 | HE OFFERED PROOF IN THE FORM OF A LARGE CHART |
| 7 | OATS ARE A FOOD EATEN BY HORSE AND MAN |
| 8 | THERE ARE MANY WAYS TO DO THESE THINGS |
| 9 | THEY SANG THE SAME TUNES AT EACH PARTY |
| 10 | THE ROOF SHOULD BE TILTED AT A SHARP SLANT |

Table S2: Complete set of sentences used in the task.

IEEE Recommended Practice for Speech Quality Measurements. (1969). *IEEE No 297-1969*, 1–24. IEEE No 297-1969. https://doi.org/10.1109/IEEESTD.1969.7405210.
